# Tools for intersectional optical and chemical tagging on cell surfaces

**DOI:** 10.1101/2024.10.29.620873

**Authors:** Sarah Innes-Gold, Hanzeng Cheng, Luping Liu, Adam E. Cohen

## Abstract

We present versatile tools for intersectional optical and chemical tagging of live cells. Photocaged tetrazines serve as “photo-click” adapters between recognition groups on the cell surface and diverse chemical payloads. We describe two new functionalized photocaged tetrazine structures which add a light-gating step to three common cell-targeting chemical methods: HaloTag/chloroalkane labeling, nonspecific primary amine labeling, and antibody labeling. We demonstrate light-gated versions of these three techniques in live cultured cells. We then explore two applications: monitoring tissue flows on the surface of developing zebrafish embryos, and combinatorial multicolor labeling and sorting of optically defined groups of cells. Photo-click adapters add optical control to cell tagging schemes, with modularity in both tag and cell attachment chemistry.

## Introduction

Light can control chemical reactions with high resolution in space and time. This capability is particularly useful in biological systems, as a means to probe spatial and temporal aspects of biological signaling. There are many methods to photo-tag cells: we and others have developed methods to select cells based on visually discernable traits by photo-crosslinking fluorescent probes to optically targeted cells of interest [1,2,3,4]. More recently, several groups have developed methods of photo-tagging cells with oligonucleotides, coupling information about their location to genomics data obtained in a later sequencing step [5,6,7].

Our goal is to add a light-gating element into several commonly used non-light-dependent live cell targeting strategies, with modularity in the cell attachment chemistry and the chemical payload. To accomplish this goal, we use photocaged dihydrotetrazine (pcDTz) [8] as a “photo-click adaptor.” The caged dihydrotetrazine is uncaged by visible light (405 nm), whereafter it can react quickly via a bio-orthogonal reaction with strained alkenes, including *trans*-cyclooctene (TCO) [9] (Figure 1A). This strategy enables fast, bio-orthogonal, light-controlled chemistry on the surface of live cells. Reference [8] presented the first of these compounds and demonstrated light-gated fluorescent labeling of an individual cell in which a lipid-conjugated pcDTz was incorporated into the membrane.

**Figure 1.**
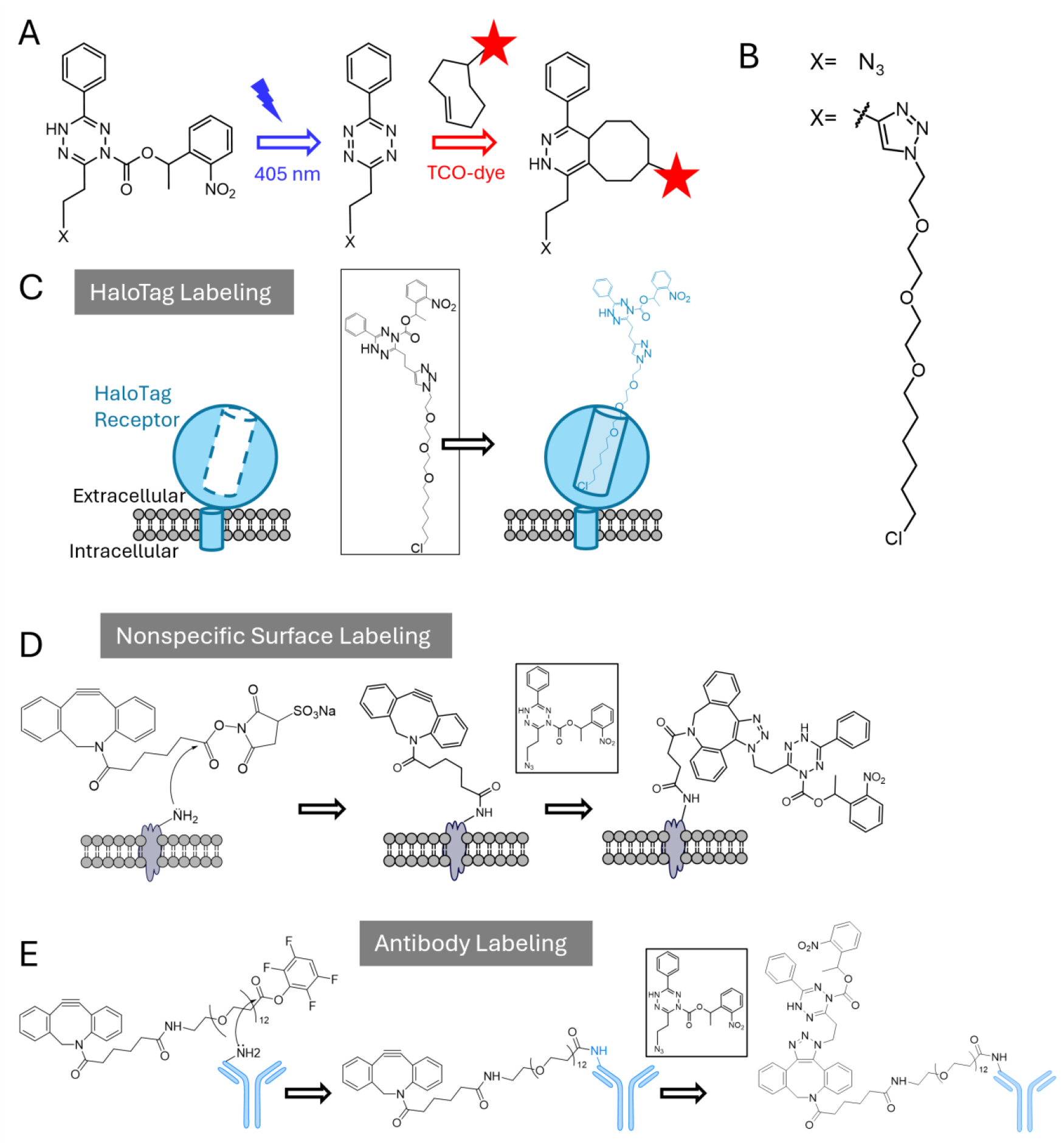
Cell-targeted photocaged tetrazines. A. Illumination of photocaged dihydrotetrazine (pcDTz) uncages it to form active tetrazine, which then reacts with TCO-conjugated dye. X denotes functional group. B. Functional groups azide and chloroalkane “HaloTag” ligand where attached to pcDTz. C. Schematic showing attachment of chloroalkane-conjugated pcDTz to a membrane-expressed HaloTag receptor. D. Schematic showing two-step attachment of pcDTz to membrane proteins via reactive ester-amine coupling and azide-DBCO click chemistry. E. Schematic showing two-step attachment of pcDTz to antibody via reactive ester-amine coupling and azide-DBCO click chemistry.

We leverage pcDTz as a photo-click adaptor in three commonly used cell targeting strategies. First, we couple light-control to the Halotag-chloroalkane system [10] for intersectional genetic and optical targeting. Second, we use nonspecific surface protein targeting via reactive esters for wide-area optical targeting [11]. Third, we incorporate the pcDTz onto antibodies for intersectional immunohistochemical and optical labeling. By adding light control to these cell-targeting strategies, we can follow the motion of tagged components; and we can increase labeling specificity by optically targeting specific morphological or functional features of the sample. To add optical control to these three methods, we use two new functional pcDTz structures. We test the structures and cell-targeting strategies in live cultured cells, demonstrating multicolor labeling at both large and subcellular scale. We explore two applications: tracking tissue flows in a developing zebrafish embryo, and combinatorial labeling for live cell sorting.

## Results

### HaloTag mediated photo-click in cultured cells

HaloTag is a powerful genetically encoded tool for tagging proteins of interest. The HaloTag receptor (HTR) forms a covalent bond with exogenous labels conjugated to a chloroalkane “HaloTag ligand” (HTL). With pcDTz fused to an HTL (Figure 1B), we added light-gating to the HaloTag toolbox.

As a demonstration, we expressed the HTR fused to the extracellular side of a transmembrane protein (platelet-derived growth factor receptor beta (PDGFR) transmembrane domain) [12]. We incubated the cells with a mixture of HTL-pcDTz and HTL-AF488, with the latter added at a small mole fraction to mark HTR-PDGFR-expressing cells. After rinsing out unbound HTL reagents, we uncaged the bound HTL-pcDTz with patterned 405 nm light, and added a TCO-conjugated fluorescent dye (Figure 1C). Figure 2A shows iterative photo-uncaging and TCO-dye labeling steps on subcellular regions (∼10 µm) of an HTR-PDGFR-expressing MDCK cell. We confirmed that TCO-dye only bound to cells (1) expressing HTR-PDGFR, as confirmed by the co-stain with HTL-AF488 AND (2) within the illuminated region (Figure S1).

**Figure 2.**
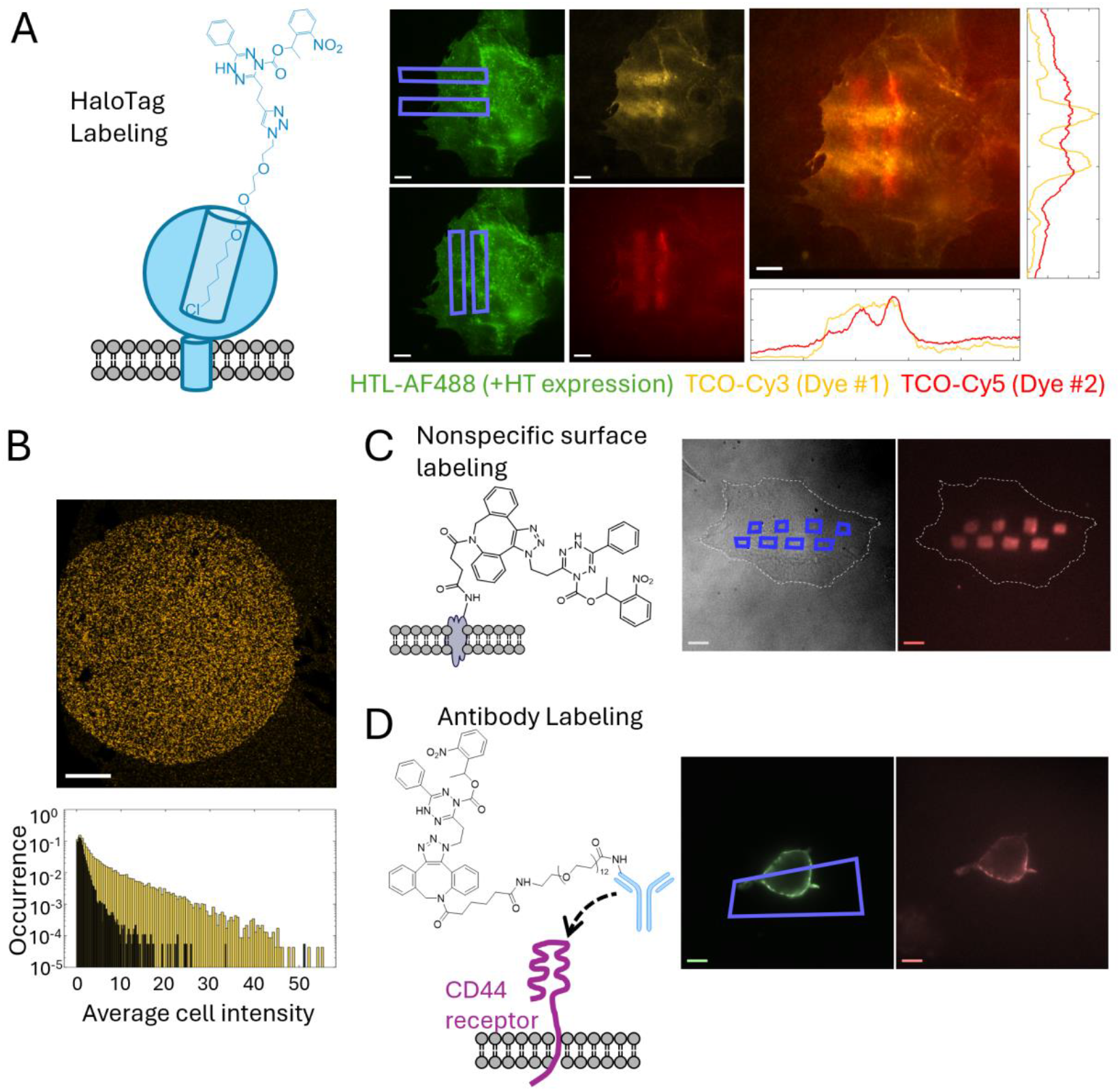
Opto-chemical tagging of cultured cells. A. Genetically targeted surface labeling. Fluorescence images showing subcellular multicolor light-gated labeling via HTL-pcDTz on an HTR-PDGFR-expressing MDCK cell. Cell was simultaneously labeled with low-concentration HTL-AF488 to indicate HTR expression (green). Cell was then illuminated with two horizontal bars of 405 nm uncaging light (blue rectangles; top left), and stained with TCO-Cy3 (yellow; top middle). Cell was then illuminated with vertical bars of 405 nm light (bottom left) and stained with TCO-Cy5 (red; bottom middle). Rightmost image merges the yellow and red fluorescence channels and shows line profile of Cy3 (yellow) and Cy5 (red) intensity across the patterned regions. Scale bar 10 µm. B. Top: Fluorescence image of a confluent layer of HEK cells expressing HTR-PDGFR and treated with HTL-pcDTz. The pcDTz was photo-uncaged in a 5 mm diameter circle and then labeled with TCO-Cy3. Scale bar 1 mm. Bottom: histogram of cell average intensities inside (yellow) and outside (black) patterned region. C. Nonspecific surface labeling. Brightfield (left) and fluorescence (right) images of a wild-type MDCK cell after treating with sulfoNHS-DBCO and azide-pcDTz, illuminating with patterned 405 nm light (purple regions, left), and perfusion with TCO-Cy5 dye (red). White dashed line shows cell outline. Scale bar 10 µm. D. Antibody-targeted surface labeling. HEK cells were transfected with CD44-HaloTag, treated with biotinylated antiCD44-pcDTz and photopatterned. Purple indicates regions targeted with 405 nm light. Left image shows fluorescence from streptavidin-AF488 (green), indicating binding of the antibody. Right image show fluorescence after uncaging and perfusion with TCO-Cy5 (red). Scale bar 10 µm.

Figure 2B demonstrates the same technique on a larger scale. We patterned a 5 mm circular region on a confluent layer of HTR-PDGFR-expressing HEK cells. Figure S2a shows the dependence of labeling density on dose of 405 nm uncaging light. The optical dose to uncage 50% of the HTL-pcDTz molecules was 198 ± 20 J/cm^2^ (mean ± s.e.m., *n* = 5 experiments). Fig. S2b shows dependence of labeling density on TCO-dye exposure. At a concentration of 10 µM, labeling reached 50% of saturation in 451 ± 22 s (mean ± s.e.m., *n* = 4 experiments). These results show that HTL-pcDTz combines the genetic specificity of the HaloTag system with the spatial specificity of photo-click.

### Nonspecific protein targeted photo-click in cultured cells

Nonspecific staining of cell surface proteins is often accomplished using *N*-hydroxysuccinimide (NHS) ester or other reactive esters, which are very reactive to primary amines. We describe a method of introducing light-gating to this useful protein-targeting approach. NHS ester groups hydrolyze readily, precluding long term storage, and complicating the synthesis. We opted instead to functionalize our pcDTz with a second click handle, azide, and attached the labile ester at the time of the experiment, using a commercially available linker.

We first used a sulfonated NHS (sulfoNHS) ester to attach the strained alkyne dibenzocyclooctyne (DBCO) to amines on the extracellular surface (the sulfonate charge prevents membrane permeation). We then added azide-pcDTz (Figure 1B), which reacted with DBCO via copper-free strain-promoted azide-alkyne cycloaddition [13]. We uncaged the pcDTz with patterned 405 nm light, and exposed the region to TCO-dyes (Figure 1D). This approach achieved subcellular labeling via nonspecific protein staining (Figure 2C).

Due to the non-specific nature of the sulfoNHS labeling, this strategy also labels proteins on the substrate surface. Thus, this approach provides a general strategy for photochemical patterning of surface chemistry. In fluorescent cell-labeling applications, one must keep in mind that the label is attached to the substrate below the cell in addition to the cell surface.

### Antibody mediated photo-click in cultured cells

Live cell immuno-targeting uses labeled antibodies to target cells that express a particular antigen. The high biomolecular specificity of antibodies makes the strategy useful for biological imaging, as well as in biotechnology and drug delivery. A primary advantage of antibody-based labeling is that the target cells do not need to be genetically modified, so the technique is compatible with primary tissues and even use in humans. For example, antibody-drug conjugates are widely used in cancer therapies [14]. However, one may wish to target a label based on the conjunction of an epitope and some other visually identifiable feature; or to target in a temporally or spatially restricted pattern. We introduce photo-control to enable these features.

We labeled primary amines on antibodies by treating first with tetrafluorophenyl ester-DBCO (plus a polyethylene glycol linker, to improve solubility) and then with azide-pcDTz (Methods). We applied this procedure to commercially available biotinylated anti-human CD44 (Figure 1E). CD44 is a membrane receptor involved in cell adhesion, growth, motility, and metastasis [15]. We expressed a CD44-HaloTag fusion [16] in HEK cells, which have very low endogenous expression of CD44 (Figure S3A). We stained the intracellularly expressed HaloTag with cell-permeant HTL-JF635 to confirm expression and membrane trafficking. In the same sample, we confirmed extracellularly accessible membrane-localized CD44 by staining with biotinylated anti-CD44 and streptavidin-AF488 (Figure S3A). We confirmed that the pcDTz-modified antibody bound to cells expressing CD44 (as indicated by streptavidin-AF488 staining) and did not bind to cells that did not express CD44 (Fig S3B, C).

We demonstrated antibody mediated photo-tagging of a subcellular region of a CD44-expressing HEK cell (Figure 2D). Both light and CD44 expression were required for labeling (Figure S3C). Successive rounds of illumination and exposure to different TCO-dyes enabled multicolor tagging (Figure S3C). As in the HaloTag system, CD44 expression was variable between cells. To control for this variability, we either co-stained for the biotin group on the antibody using fluorescently tagged streptavidin (Figure 2D), or labeled the intracellular HaloTag (Figure S3C). PcDTZ-antibody conjugates combine the biological specificity of antibody targeting with the spatial specificity of light. We next explored applications of intersectional optical and chemical targeting.

### Photo-click on live zebrafish embryos

The spatiotemporal control, multicolor capability, and live cell compatibility of pcDTz labeling make this approach attractive for imaging flows in biology. We explored this application area by tracking flows on the surface of a live zebrafish embryo.

Zebrafish embryos develop rapidly in the first 24 hours after fertilization and are transparent during this stage [17]. Tracking all individual cells is possible [18,19], but becomes technically challenging and computationally expensive as the embryo grows, motivating the search for simple methods to track tissue flows without the need for continuous imaging. Recent efforts have explored methods to pattern a registration grid onto live tissue, for tracking or downstream omics [20]. We show that pcDTz photo-click could be similarly useful. We first functionalized the embryo surface with sulfoNHS ester-DBCO, then added azide-pcDTz. We iteratively uncaged and added TCO-dyes to pattern the embryo tail with perpendicular stripes in two colors (Figure 3A, B, D). Embryos tolerated this process well and continued to develop at a similar rate to untreated control embryos (Figure S4). We imaged the patterned embryos 12 hours after staining. The patterns were still clearly visible, but the stripes had been distorted by embryo growth, showing greater broadening on the ventral side of the tail compared to the dorsal side, as the tail straightened (Figure 3C, E). Three-dimensional reconstructions confirmed that the labeling was limited to a superficial layer, as expected for the non-cell permeant sulfoNHS ester.

**Figure 3.**
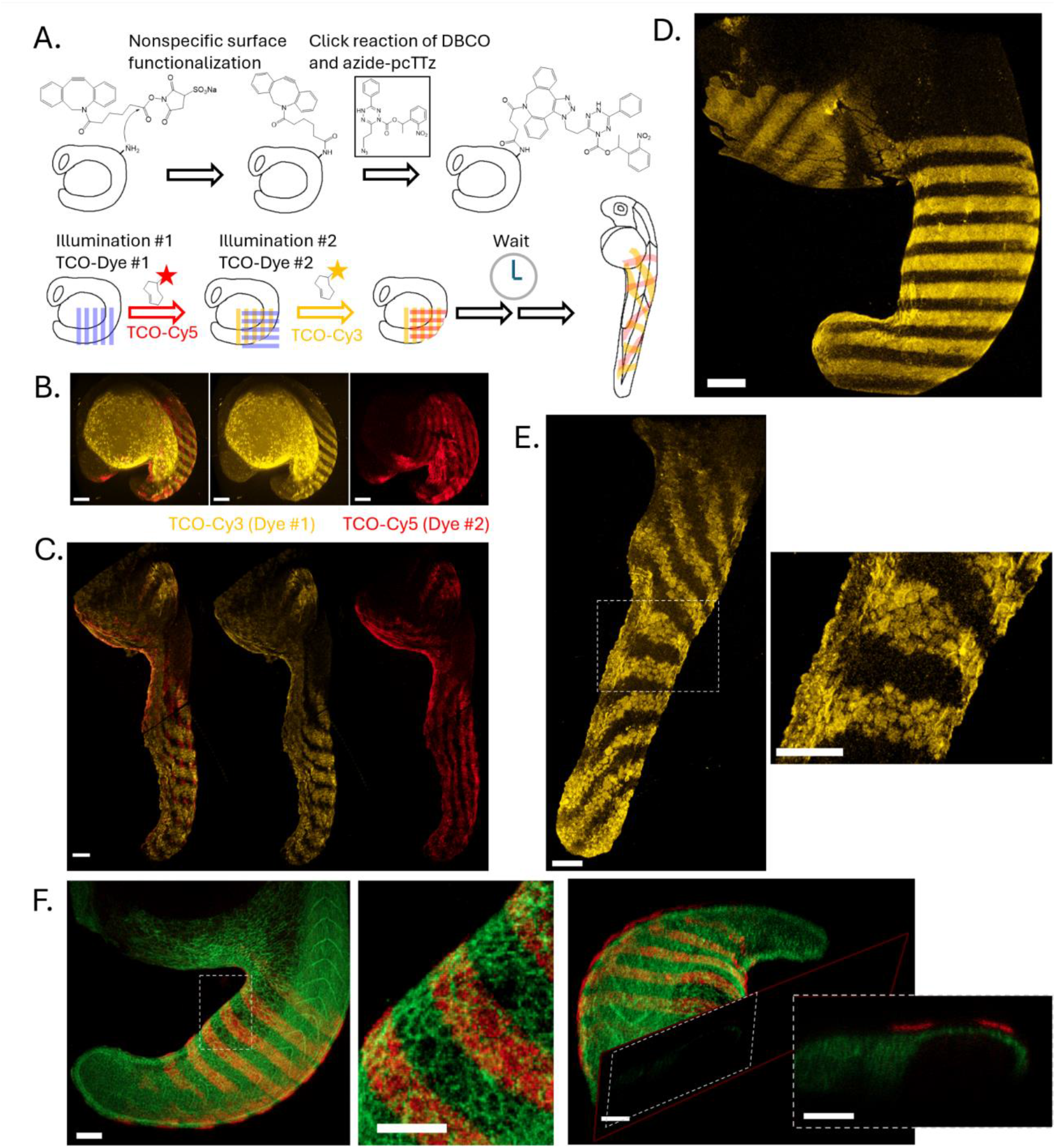
Amine-targeted photo-click on live zebrafish embryos. A. Schematic showing chemical treatment and iterative patterning of stripes on zebrafish embryo tail. B. Zebrafish embryo, 14 hours post-fertilization (hpf), treated with sulfoNHS-DBCO and azide-pcDTz, patterned iteratively with 405 nm light, TCO-Cy3 (dye #1, yellow), and TCO-Cy5 (dye #2, red). C. Zebrafish embryo that underwent same patterning process described in (B), then left for 12 hours before fixation and imaging. D. Zebrafish embryo, with Cy3 stripes patterned across the tail at ∼18 hpf, followed by immediate fixation and imaging. E. Zebrafish embryo, with Cy3 stripes patterned across the tail at ∼18 hpf, then left until 24 hpf before fixation and imaging. Inset shows boxed region, magnified. F. TCO-Cy5 was patterned on the surface of a zebrafish embryo ubiquitously expressing cell-membrane-localized mNeonGreen. Three-dimensional imaging showed that the patterning was confined to a superficial layer. Scale bars 100 µm.

We believe that multi-color photo-click patterning is a useful approach to observe surface tissue flows in development without the need for continuous imaging. This technique may also be useful in studies of wound healing or disease progression (e.g. tumor growth), where one wished to track the motion of large ensembles of cells. While we here describe nonspecific amine-targeted photo-click, one could instead use the genetically encoded HaloTag approach to track a genetically specified population of cells.

### Photo-click labeling for cell sorting

Photo-click chemistry can be used for combinatorial labeling. With *N* distinguishable dyes and a binary labeling scheme (present or absent) one can distinguish 2^N^-1 distinct populations (assuming the all-absent condition is not detectable). This may be useful for cell sorting. In principle, the labeling can be targeted based on any observable feature of the cells. Cells with distinct spectral barcodes can then be sorted into different wells for downstream characterization, e.g. via sequencing, proteomics, or other biochemical analyses. The ability to sort cells based on arbitrary features into multiple buckets would be a powerful enabling capability for optical pooled screens or for characterizing heterogeneous cell populations [21–25].

Here, we used three TCO-dyes (Cy3, Cy5, and AZDye488), to distinguishably label seven subpopulations of cells within a culture dish. We attached pcDTz to a near-confluent layer of live HEK cells (either via sulfoNHS-DBCO + azide-pcDTz, Figure 4, or via HTR-PDGFR expression + HTL-pcDTz, Figure S5). We iteratively uncaged the pcDTz in partially overlapping circular regions, and incubated with TCO-dyes. For the uncaging step we used a light dose of 50 J/cm^2^, well below saturation, so that caged pcDTz remained available for subsequent uncaging steps. For the labeling, we used a dose of 10 µM, 12 minutes, which came close to saturating the labeling reaction, so that subsequent labeling steps would not follow prior uncaging patterns. Figures 4 and S5 show the resulting cells on the surface. We then dissociated the labeled cells and separated them with a fluorescence-activated cell sorter (FACS, Figure 4B). We detected clearly distinct populations corresponding to each of the seven labeling conditions (Figure 4C). These data demonstrate combinatorial labeling with pcDTz photo-click chemistry.

**Figure 4.**
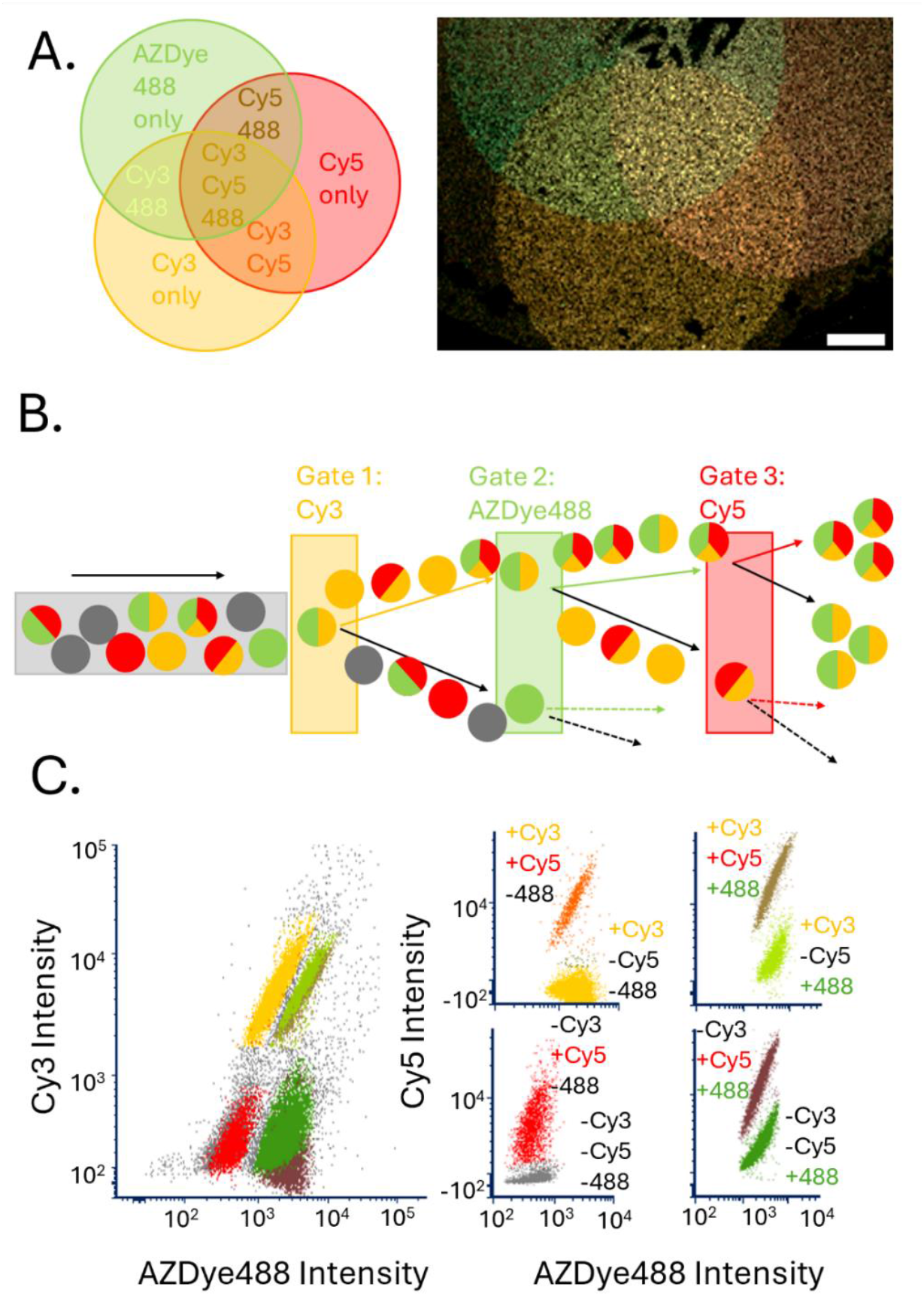
Multicolor combinatorial cell sorting. A. Left: schematic showing expected dye retention in each iteratively patterned region. Right: merged fluorescence image and schematic showing regions of HEK cells photo-patterned in partially overlapping regions by sequential targeted illumination and TCO-dye labeling (TCO-Cy3, TCO-Cy5, TCO-AZDye488). Scale bar 1 mm. B. Schematic showing FACS-sorted cell populations after patterning via sulfoNHS-DBCO + azide-pcDTz. C. FACS data showed separable populations with all seven combinations of the three dyes.

Figure S5 shows results from a similar combinatorial labeling approach in cells expressing HTR-PGDFR, with cells categorized in the dish by image analysis. Due to cell-to-cell variation in expression levels, we did not obtain entirely separable populations, either with FACS or in-dish imaging. We could, however, establish a cell intensity threshold which separated patterned and HaloTag-expressing cells from the dark cells (Figure S5). Presumably, in a monoclonal cell line expressing HT-PGDFR the expression would be more uniform across cells.

With careful control of illumination dose and labeling rates, this technique might be extended to discern multiple levels of each dye (rather than the binary case shown above), expanding the number of selectable groups without the need for additional fluorophores. Combinatorial labeling of antibody-pcDTz-labeled cells is presumably possible too.

## Discussion

Previous work established that pcDTz leverages the advantages of click chemistry (fast, easy, biocompatible), light-gated reactions (spatial control), and modular probes. Here, we also enable intersectional levels of control by combining pcDTz with chemically and genetically specific cell attachment strategies. We present two new pcDTz structures, chloroalkane “HaloTag ligand”-pcDTz and click-capable azide-pcDTz. With these structures, we add light-control capabilities to three popular cell-attachment strategies: HaloTag, NHS ester amine chemistry, and antibody labeling. We explore two potential applications for the technique: tracking tissues flows on the surface of the live zebrafish embryo, and multicolor, combinatorial live cell labeling and sorting.

We foresee a range of future directions for pcDTz photo-click. A promising application is in tracking dynamics of proteins within cells. Here, we demonstrated tracking of tissue flows in embryos on a millimeter scale, with nonspecific protein labeling. Light control also enables patterns to be targeted subcellularly (<10 µm), and targeted to specific proteins via a HaloTag linker. Future work will use these approaches to track the motion of specific protein sub-populations within cells. Superresolution imaging techniques may enable very high resolution tracking.

In a separate area, we envision the cell-sorting application explored in the previous section will be useful for optical phenotypic screening, and combined with subsequent omics. In expanding the scope of live cell and biomedically motivated applications, we also plan to explore red-shifted varieties of pcDTz [8], for deeper tissue penetration and reduced phototoxicity.

## Methods

*Synthesis*: The functionalized pcDTz structures used here were prepared from 1-(2-nitrophenyl)ethyl 6-(but-3-yn-1-yl)-3-phenyl-1,2,4,5-tetrazine-1(4*H*)-carboxylate and 2-(6-phenyl-1,2,4,5-tetrazin-3-yl)ethan-1-ol, whose syntheses are described previously [8,26].

Procedure to prepare 1-(2-nitrophenyl)ethyl6-(2-(1-(2-(2-(2-((6chlorohexyl) oxy)ethoxy)ethoxy)ethyl)-1H-1,2,3-triazol-4-yl)ethyl)-3-phenyl-1,2,4,5-tetrazine-1(4*H*) - carboxylate (HTL-pcDTz):

Under argon, 12 mL of CH_3_CN/H_2_O (v/v = 2/1) was added to a mixture of photocaged dihydrotetrazine 1-(2-nitrophenyl)ethyl 6-(but-3-yn-1-yl)-3-phenyl-1,2,4,5-tetrazine-1(4*H*)-carboxylate (127 mg, 0.31 mmol), 1-(2-(2-(2-azidoethoxy)ethoxy)ethoxy)-6-chlorohexane (125 mg, 0.42 mmol), CuI (106 mg, 0.56 mmol), and sodium L-ascorbate (110 mg, 0.56 mmol) at room temperature. The reaction mixture was stirred at 37 °C, overnight. Upon completion, the reaction solvent was removed under reduced pressure. The residue was purified by column chromatography (4% MeOH/CH_2_Cl_2_), and the desired product HLT-pcDTz was obtained (71 mg, 32% yield).

Procedure to prepare 1-(2-nitrophenyl)ethyl 6-(2-azidoethyl)-3-phenyl-1,2,4,5-tetrazine-1(4*H*)-carboxylate (azide-pcDTz):

Under argon, triethylamine (617 μL, 4.5 mmol), MsCl (343 μL, 4.5 mmol) and DMAP (182 mg, 1.5 mmol) were added to a solution of 2-(6-phenyl-1,2,4,5-tetrazin-3-yl)ethan-1-ol (300 mg, 1.5 mmol) in CH_2_Cl_2_ (8 mL) at room temperature. Upon completion, the reaction mixture was concentrated under reduced pressure. The residue was purified by column chromatography (CH_2_Cl_2_), and the 2-(6-phenyl-1,2,4,5-tetrazin-3-yl)ethyl methanesulfonate was obtained. Then, NaN_3_ (309 mg, 4.75 mmol) was added to a solution of 2-(6-phenyl-1,2,4,5-tetrazin-3-yl)ethyl methanesulfonate in DMF (6 mL), the mixture stirred at room temperature under argon for 6 h. Then, the reaction mixture was extracted with CH_2_Cl_2_ and washed with a saturated solution of NaCl in water. The extract was combined, dried over Na_2_SO_4_, and concentrated under reduced pressure. The residue was purified by column chromatography (50% Hexane/EtOAc), and 3-(2-azidoethyl)-6-phenyl-1,2,4,5-tetrazine was obtained (132 mg, 40% yield). Under argon, the thiourea dioxide (76 mg, 0.7 mmol) was added to a solution of 3-(2-azidoethyl)-6-phenyl-1,2,4,5-tetrazine (80 mg 0.35 mmol) in 15 mL of DMF/H_2_O (v/v = 1/2) at room temperature. The reaction mixture was stirred at 95 °C for 1-2 hours. Upon completion, the color of the reaction mixture changed from pink to light yellow. Under argon, 50 mL of EtOAc was added the reaction mixture, which was washed by 30 mL of H_2_O. The extract was combined and concentrated by reduced pressure. The residue was purified by column chromatography (CH_2_Cl_2_ as the eluent), and 3-(2-azidoethyl)-6-phenyl-1,4-dihydro-1,2,4,5-tetrazine was obtained (60 mg, 74% yield). Under argon, the solution of 1-(2-nitrophenyl)ethyl carbonochloridate (158 mg, 0.65 mmol) in toluene (1 mL) was added to a solution of 3-(2-azidoethyl)-6-phenyl-1,4-dihydro-1,2,4,5-tetrazine (60 mg 0.26 mmol) in pyridine (5 mL) at 0 °C. The reaction mixture was stirred at 37 °C for 24 h. Upon completion, the reaction solvent was removed under reduced pressure. The residue was purified by column chromatography (1% MeOH/ CH_2_Cl_2_), and the azide-pcDTz was obtained (70 mg, 83% yield).

Full experimental details and characterization of the newly described compounds (^1^H NMR, ^13^C NMR, mass spectra) are provided in the Supporting Information.

### Cell culture and transfection

HEK293T and MDCK cells (ATCC) were cultured in Dulbecco’s modified Eagle medium (DMEM) supplemented with 10% fetal bovine serum, 1% GlutaMax-I, penicillin (100 U/ml), and streptomycin (100 µg/ml). Cells were transfected 24 h after passage using TransIT-293 transfection reagents (Mirus Bio) for the HEK cells, and TransIt-X2 transfection reagents (Mirus Bio) for MDCK cells. Experiments on expressing cells were performed 24 h after transfection.

#### Plasmids used

pcDNA5/FRT/TO_IgKchL_HA_HaloTag9_myc_PDGFRtmb (for expression of HaloTag on the outer membrane) was a gift from Kai Johnsson (Addgene plasmid # 175544 ; RRID:Addgene_175544) [12], and pEGFPN1-CD44-SGx3-Halo7 (for expression of CD44 with an intracellular HaloTag) was generously shared by Tim Yeh [16]. Plasmids were diluted 1:1 and 1:5 respectively in puc19 vector (New England Biolabs). HaloTag expression was confirmed using HaloTag ligand dyes HTL-AF488, HTL-JF635 (Promega CS3495B06, HT1050), 50 nM in culture medium, 30 min, followed by rinsing and, for membrane permeant JF635, another 30 min incubation in fresh media. CD44 expression was confirmed using CD44 Monoclonal Antibody (IM7)-Biotin, eBioscience (Life Technologies, 13-0441-82) and Streptavidin-AlexaFluor 488 conjugate (Life Technologies, S11223).

### Cell surface chemistry

HaloTag-expressing cells were incubated with HTL-pcDTz, 1 µM in 0.1% DMSO in culture medium, 30 min, rinsed 3x, and incubated again in fresh medium for 30 min. For nonspecific surface labeling experiments, sulfonated *N*-hydroxysuccinimide-DBCO (Click Chemistry Tools) was dissolved in anhydrous DMSO and added to cells, 100 µM in 0.1% DMSO in PBS, 10 min, then rinsed 3x. Azide-pcDTz was then added to cells at 1 µM in 0.1% DMSO in culture medium, incubated 60 min, rinsed 3x, and incubated 30 more min in fresh medium. For antibody experiments, cells were first blocked for 60 min with 3% bovine serum albumin (BSA) in Hank’s buffered salt solution (HBSS). Functionalized antibody was added at ∼10 nM in HBSS + 1% BSA for 30 min. Dish was rinsed 3x, then stained with HTL dye or streptavidin-AF488. All incubation steps were performed at 37 °C, 5% CO_2_, and all media and cell buffers were pre-warmed to 37 °C.

### Preparation of functionalized antibody

CD44 Monoclonal Antibody (IM7), Biotin, eBioscience (Life Technologies, 13-0441-82) was used after solvent exchange to PBS, pH 7.2, using BSA-passivated Amicon Ultra 10 kDa spin columns to remove sodium azide. EZ-Link TFP Ester-PEG_12_-DBCO (Life Technologies), dissolved in DMSO, was added to the antibody in ∼50x molar excess. The reaction proceeded at room temperature for 1 hour on a shaker, and overnight at 4 °C. The reaction was quenched with Tris HCl buffer, then de-salted again on a freshly BSA-passivated 10 kDa spin column to remove unreacted DBCO reagent. Antibody concentration and degree of DBCO labeling were estimated using a NanoDrop spectrophotometer, finding recovered [antibody] ∼ 1-2 µM and degree of DBCO labeling ∼ 0.7. Azide-pcDTz, dissolved in DMSO, was then added in ∼10x molar excess over the DBCO, and reacted at room temperature for 1 hour on a shaker, then overnight at 4 °C. The resulting product was diluted 1:100 in HBSS immediately before addition to cells. Functionalized antibody was stored at 4 °C and used within five days.

### Zebrafish embryo surface chemistry

All vertebrate experiments were approved by the Institutional Animal Care and Use Committee of Harvard University (protocols 10-13-4 and 21-06-391-1). The zebrafish (*Danio rerio*) AB wild-type strain was used for all experiments. Adult fish were raised at 28.5 °C on a 14 h light/10 h dark cycle. Embryos were collected by crossing female and male adults (3–24 months old). Embryos were collected and dechorionated at approximately 10 hours post fertilization (hpf) by immersing in 1 mg*/*ml pronase protease (Sigma) in 0.3× Danieau’s buffer (17.4 mM NaCl, 0.21 mM KCl, 0.12 mM MgSO_4_, 0.18 mM Ca(NO_3_) _2_, 1.5 mM HEPES, pH 7.2) for 7 min at room temperature. Embryos were then treated with 100 µM sulfoNHS-DBCO for 10 min at room temperature, rinsed, and treated with 10 µM azide-pcDTz for 30-60 min at room temperature. They were then rinsed and incubated in Danieu’s buffer for 30-120 minutes at room temperature before photo-uncaging experiments.

### Photo-uncaging

Patterning of individual cells or of fish embryos was performed on a home-made inverted epifluorescence microscope. A 405 nm laser (Lasever, LSR405NL-200, 200 mW) was patterned by a digital micromirror device (Texas Instruments, DLP3000) and re-imaged onto the sample via a 60x water immersion objective (Olympus, UPLSAPO60XW). Illumination dose was ∼500 J/cm^2^ for near complete uncaging.

Cells were plated on Matrigel or poly-D-lysine-coated glass, and immersed in an “imaging buffer” containing (in mM): 125 NaCl, 2.5 KCl, 2 CaCl_2_, 1 MgCl_2_, 15 HEPES, 25 glucose (pH 7.3). Individual cells were perfused with TCO-dye using a BioPen single cell application system (Fluicell). TCO-dyes used were TCO-Cy3 (AAT Bioquest), TCO-Cy5 (Click Chemistry Tools), TCO-AZDye488 (Click Chemistry Tools), and TCO-CF640R (Biotium).

Fish embryos were placed in wells on an agarose-coated glass dish for patterning. Fish embryos were transferred to a separate dish for TCO-dye immersion and rinsing.

Wide-area patterning of confluent cell layers was performed on a wide-area upright microscope, similar to the “Firefly” design described previously [27]. Light from a 405 nm laser (Changchun New Industries, PSU-H-LED, 1W) was patterned by a digital micromirror device (Vialux V7000). Samples were illuminated at a dose of ∼50 J/cm^2^ for each round of uncaging. TCO-dye and rinsing steps were performed by hand pipetting.

### Microscopy

Patterning and single-cell imaging were performed on a custom-built inverted epifluorescence microscope. Zebrafish embryos were fixed in 4% paraformaldehyde (Thermo Fisher) before the final imaging step. Zebrafish embryo confocal z-stacks were obtained at the Harvard Center for Biological Imaging, at 10 and 20x using a LSM 980 NLO Multi-Photon Microscope (Zeiss). Wide-area cell images were obtained at the Harvard Center for Biological Imaging, at 10 and 20x using a LSM 900 Live Cell Confocal Microscope (Zeiss).

### Flow cytometry

After patterning, cells were dissociated by treatment with trypsin (Thermo Fisher, 15050057), quenched with culture medium, spun down (300 g, 5 min), and resuspended in imaging buffer supplemented with 2% fetal bovine serum. Flow cytometry and FACS were performed at the Flow Cytometry Core at the Harvard University Bauer Core Facility, using a BD FACS Aria Fusion Cell Sorter with two rounds of 4-way sorting, and laser lines at 488 nm, 561 nm, and 637 nm, for TCO-AZDye488, TCO-Cy3, and TCO-Cy5 dye.

### Image analysis

Image processing of confocal z-stacks and wide-area cell images was performed in Zen Blue software (Zeiss). Single cell images were analyzed in MATLAB (Mathworks). Cell segmentation for classification from image (Figure S5) was performed using Ilastik: cells were segmented based on a nuclear DAPI stain, and a mask for each cell was exported to MATLAB. Masks were expanded to include the labeled membrane, and average intensity was calculated for each cell to generate cell intensity histograms.

## Supporting information

Supp. Figures and Synthesis details

## Acknowledgments

We thank Andrew Preecha, Shahinoor Begum, and Camila Bodden for technical assistance. We thank Bill Jia for help with fish embryos and Madeleine Howell, Yitong Qi, and Daniel Itkis for help with the microscopy setups. We thank Tim (Hsin-Chih) Yeh for the CD44-HaloTag plasmid. We thank Heather Brown-Harding, Alex Lovely, and Douglas Richardson at the Harvard Center for Biological Imaging. We thank Jeffery Nelson at the Bauer Flow Cytometry core for help with FACS experiments. We thank Jessica Miller and Karen Hurley at the Zebrafish Facility. This work was supported by Schmidt Futures and the Gordon and Betty Moore Foundation, and the Nonprofit Central Research Institute Fund of Chinese Academy of Medical Sciences (2022-RC350-02), CAMS Innovation Fund for Medical Sciences (2021-I2M-1-026 and 2023-I2M-2-006), National Natural Science Foundation of China (Nos. 82404404), and Beijing Nova Program to L. L. (No. 2022048). S.I-G was supported by a postdoctoral fellowship from the Jane Coffin Childs Memorial Fund for Medical Research.

## Supporting Information

Supplementary Figures 1-5, details of chemical synthesis methods, and NMR and mass spec characterization of new compounds.

## References

(1) Chien, M.-P.; Werley, C. A.; Farhi, S. L.; Cohen, A. E. Photostick: A Method for Selective Isolation of Target Cells from Culture. Chem. Sci. 2015, 6 (3), 1701–1705. 10.1039/C4SC03676J.

(2) Binan, L.; Mazzaferri, J.; Choquet, K.; Lorenzo, L.-E.; Wang, Y. C.; Affar, E. B.; De Koninck, Y.; Ragoussis, J.; Kleinman, C. L.; Costantino, S. Live Single-Cell Laser Tag. Nat. Commun. 2016, 7 (1), 11636. 10.1038/ncomms11636.

(3) Kuo, C.-T.; Thompson, A. M.; Gallina, M. E.; Ye, F.; Johnson, E. S.; Sun, W.; Zhao, M.; Yu, J.; Wu, I.-C.; Fujimoto, B.; DuFort, C. C.; Carlson, M. A.; Hingorani, S. R.; Paguirigan, A. L.; Radich, J. P.; Chiu, D. T. Optical Painting and Fluorescence Activated Sorting of Single Adherent Cells Labelled with Photoswitchable Pdots. Nat. Commun. 2016, 7 (1), 11468. 10.1038/ncomms11468.

(4) Binan, L.; Bélanger, F.; Uriarte, M.; Lemay, J. F.; Pelletier De Koninck, J. C.; Roy, J.; Affar, E. B.; Drobetsky, E.; Wurtele, H.; Costantino, S. Opto-Magnetic Capture of Individual Cells Based on Visual Phenotypes. eLife 2019, 8, e45239. 10.7554/eLife.45239.

(5) Hu, K. H.; Eichorst, J. P.; McGinnis, C. S.; Patterson, D. M.; Chow, E. D.; Kersten, K.; Jameson, S. C.; Gartner, Z. J.; Rao, A. A.; Krummel, M. F. ZipSeq: Barcoding for Real-Time Mapping of Single Cell Transcriptomes. Nat. Methods 2020, 17 (8), 833–843. 10.1038/s41592-020-0880-2.

(6) Kishi, J. Y.; Liu, N.; West, E. R.; Sheng, K.; Jordanides, J. J.; Serrata, M.; Cepko, C. L.; Saka, S. K.; Yin, P. Light-Seq: Light-Directed in Situ Barcoding of Biomolecules in Fixed Cells and Tissues for Spatially Indexed Sequencing. Nat. Methods 2022, 19 (11), 1393–1402. 10.1038/s41592-022-01604-1.

(7) Mangiameli, S. M.; Chen, H.; Earl, A. S.; Dobkin, J. A.; Lesman, D.; Buenrostro, J. D.; Chen, F. Photoselective Sequencing: Microscopically Guided Genomic Measurements with Subcellular Resolution. Nat. Methods 2023, 20 (5), 686–694. 10.1038/s41592-023-01845-8.

(8) Liu, L.; Zhang, D.; Johnson, M.; Devaraj, N. K. Light-Activated Tetrazines Enable Precision Live-Cell Bioorthogonal Chemistry. Nat. Chem. 2022, 1–8. 10.1038/s41557-022-00963-8.

(9) Blackman, M. L.; Royzen, M.; Fox, J. M. Tetrazine Ligation: Fast Bioconjugation Based on Inverse-Electron-Demand Diels–Alder Reactivity. J. Am. Chem. Soc. 2008, 130 (41), 13518–13519. 10.1021/ja8053805.

(10) Los, G. V.; Encell, L. P.; McDougall, M. G.; Hartzell, D. D.; Karassina, N.; Zimprich, C.; Wood, M. G.; Learish, R.; Ohana, R. F.; Urh, M.; Simpson, D.; Mendez, J.; Zimmerman, K.; Otto, P.; Vidugiris, G.; Zhu, J.; Darzins, A.; Klaubert, D. H.; Bulleit, R. F.; Wood, K. V. HaloTag: A Novel Protein Labeling Technology for Cell Imaging and Protein Analysis. ACS Chem. Biol. 2008, 3 (6), 373–382. 10/dsvwdc.

(11) Mattson, G.; Conklin, E.; Desai, S.; Nielander, G.; Savage, M. D.; Morgensen, S. A Practical Approach to Crosslinking. Mol. Biol. Rep. 1993, 17 (3), 167–183. 10.1007/BF00986726.

(12) Frei, M. S.; Tarnawski, M.; Roberti, M. J.; Koch, B.; Hiblot, J.; Johnsson, K. Engineered HaloTag Variants for Fluorescence Lifetime Multiplexing. Nat. Methods 2022, 19 (1), 65–70. 10.1038/s41592-021-01341-x.

(13) Agard, N. J.; Baskin, J. M.; Prescher, J. A.; Lo, A.; Bertozzi, C. R. A Comparative Study of Bioorthogonal Reactions with Azides. ACS Chem. Biol. 2006, 1 (10), 644–648. 10.1021/cb6003228.

(14) Beck, A.; Goetsch, L.; Dumontet, C.; Corvaïa, N. Strategies and Challenges for the next Generation of Antibody–Drug Conjugates. Nat. Rev. Drug Discov. 2017, 16 (5), 315–337. 10.1038/nrd.2016.268.

(15) Ponta, H.; Sherman, L.; Herrlich, P. A. CD44: From Adhesion Molecules to Signalling Regulators. Nat. Rev. Mol. Cell Biol. 2003, 4 (1), 33–45. 10.1038/nrm1004.

(16) Karam, J.; Singer, B. J.; Miwa, H.; Chen, L. H.; Maran, K.; Hasani, M.; Garza, S.; Onyekwere, B.; Yeh, H.-C.; Li, S.; Carlo, D. D.; Seidlits, S. K. Molecular Weight of Hyaluronic Acid Crosslinked into Biomaterial Scaffolds Affects Angiogenic Potential. Acta Biomater. 2023, 169, 228–242. 10.1016/j.actbio.2023.08.001.

(17) Kimmel, C. B.; Ballard, W. W.; Kimmel, S. R.; Ullmann, B.; Schilling, T. F. Stages of Embryonic Development of the Zebrafish. Dev. Dyn. 1995, 203 (3), 253–310. 10.1002/aja.1002030302.

(18) Keller, P. J.; Schmidt, A. D.; Wittbrodt, J.; Stelzer, E. H. K. Reconstruction of Zebrafish Early Embryonic Development by Scanned Light Sheet Microscopy. Science 2008, 322 (5904), 1065–1069. 10.1126/science.1162493.

(19) Bragantini, J.; Theodoro, I.; Zhao, X.; Huijben, T. A. P. M.; Hirata-Miyasaki, E.; VijayKumar, S.; Balasubramanian, A.; Lao, T.; Agrawal, R.; Xiao, S.; Lammerding, J.; Mehta, S.; Falcão, A. X.; Jacobo, A.; Lange, M.; Royer, L. A. Ultrack: Pushing the Limits of Cell Tracking across Biological Scales. bioRxiv September 3, 2024, p 2024.09.02.610652. 10.1101/2024.09.02.610652.

(20) Cotterell, J.; Swoger, J.; Robert-Moreno, A.; Cardona, H.; Musy, M.; Sharpe, J. Cell 3D Positioning by Optical Encoding (C3PO) and Its Application to Spatial Transcriptomics. bioRxiv March 12, 2024, p 2024.03.12.584578. 10.1101/2024.03.12.584578.

(21) Feldman, D.; Singh, A.; Schmid-Burgk, J. L.; Carlson, R. J.; Mezger, A.; Garrity, A. J.; Zhang, F.; Blainey, P. C. Optical Pooled Screens in Human Cells. Cell 2019, 179 (3), 787-799.e17. 10.1016/j.cell.2019.09.016.

(22) Hasle, N.; Cooke, A.; Srivatsan, S.; Huang, H.; Stephany, J. J.; Krieger, Z.; Jackson, D.; Tang, W.; Pendyala, S.; Monnat, R. J.; Trapnell, C.; Hatch, E. M.; Fowler, D. M. High-throughput, Microscope-based Sorting to Dissect Cellular Heterogeneity. Mol. Syst. Biol. 2020, 16 (6), e9442. 10.15252/msb.20209442.

(23) Kanfer, G.; Sarraf, S. A.; Maman, Y.; Baldwin, H.; Dominguez-Martin, E.; Johnson, K. R.; Ward, M. E.; Kampmann, M.; Lippincott-Schwartz, J.; Youle, R. J. Image-Based Pooled Whole-Genome CRISPRi Screening for Subcellular Phenotypes. J. Cell Biol. 2021, 220 (2), e202006180. 10.1083/jcb.202006180.

(24) Walton, R. T.; Singh, A.; Blainey, P. C. Pooled Genetic Screens with Image-based Profiling. Mol. Syst. Biol. 2022, 18 (11), e10768. 10.15252/msb.202110768.

(25) Tian, H.; Davis, H. C.; Wong-Campos, J. D.; Park, P.; Fan, L. Z.; Gmeiner, B.; Begum, S.; Werley, C. A.; Borja, G. B.; Upadhyay, H.; Shah, H.; Jacques, J.; Qi, Y.; Parot, V.; Deisseroth, K.; Cohen, A. E. Video-Based Pooled Screening Yields Improved Far-Red Genetically Encoded Voltage Indicators. Nat. Methods 2023, 20 (7), 1082–1094. 10.1038/s41592-022-01743-5.

(26) Mao, W.; Shi, W.; Li, J.; Su, D.; Wang, X.; Zhang, L.; Pan, L.; Wu, X.; Wu, H. Organocatalytic and Scalable Syntheses of Unsymmetrical 1,2,4,5-Tetrazines by Thiol-Containing Promotors. Angew. Chem. Int. Ed. 2019, 58 (4), 1106–1109. 10.1002/anie.201812550.

(27) Werley, C. A.; Chien, M.-P.; Cohen, A. E. Ultrawidefield Microscope for High-Speed Fluorescence Imaging and Targeted Optogenetic Stimulation. Biomed. Opt. Express 2017, 8 (12), 5794–5813. 10/gg83rc.

