## Supplementary material for "Tools for intersectional optical and chemical tagging on cell surfaces": Supp. Figures and Synthesis details

**Supporting Information**

Supplementary Figures
Chemical synthesis details
NMR and Mass Spec characterization of new compounds

**Supplementary Figures**

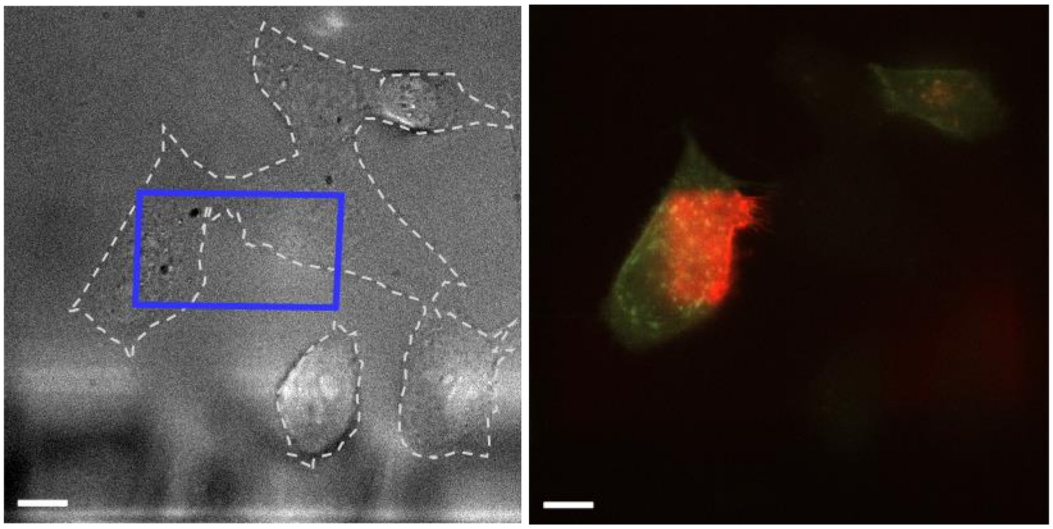

**Figure S1. Sub-cellular intersectional genetic and optical targeting.** Brightfield (left) and merged fluorescence (right) images of MDCK cells expressing HTR-PDGFR and stained with a mixture of HTL-pcDTz and HTL-AF488, uncaged in a small region with 405 nm light, and perfused with TCO-Cy5 dye. AF488 fluorescence (green) indicates HaloTag expression, Cy5 fluorescence (red) shows TCO-dye binding, gray dash line outlines all cells in brightfield image, blue line indicates region targeted with uncaging light. The pattern of Cy5 fluorescence demonstrates that the photochemical targeting only occurred in cells expressing HTR-PDGFR. The blurry objects at the bottom of the transmitted light image are the channels of the BioPen micro-pipette used for local dye perfusion. Scale bars 10 μm.

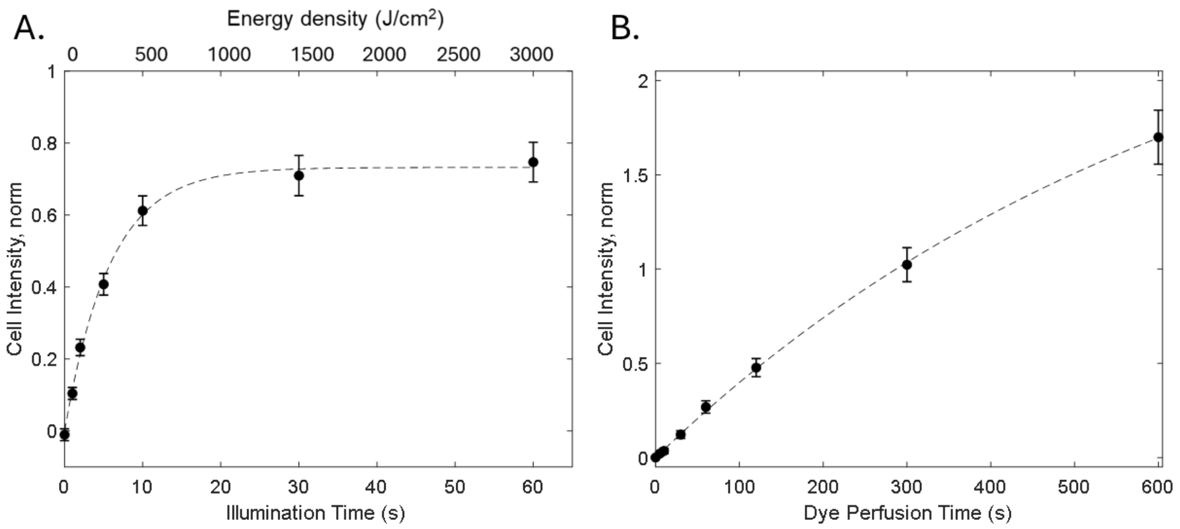


**Figure S2. Photo-click reaction kinetics.** Uncaging and binding kinetics on cells expressing HTR-PDGFR, treated with HTL-pcDTz and HTL-AF488 (10:1). Fluorescence intensity of the TCO-Cy5 signal is normalized by the HTL-AF488 signal on the same cell, to control for variations in expression level. A. Normalized Cy5 fluorescence intensity vs. illumination time (50 $W/cm^2$, 405 nm) after uncaging and 60 s perfusion with 10 μM TCO-Cy5, fit to two-parameter exponential model $I = A(1 - e^{-t/\tau})$, with characteristic timescale $\tau$ = 5.8 s [95% CI: 5.0, 6.6] and saturation value $A$ = 0.73 [95% CI: 0.70, 0.76]. N = 5 cells, room temperature. Error bars show s.e.m. B. Normalized Cy5 fluorescence intensity vs. dye perfusion time. Region of interest was first illuminated for 5 s at 125 $W/cm^2$ (625 $J/cm^2$), then perfused with 10 μM TCO-Cy5 dye. Fit to two-parameter exponential model, with characteristic timescale $\tau$ = 665 s [95% CI: 577, 752] and saturation value $A$ = 2.85 [95% CI: 2.59, 3.11]. N = 4 cells, room temperature. Error bars show s.e.m.

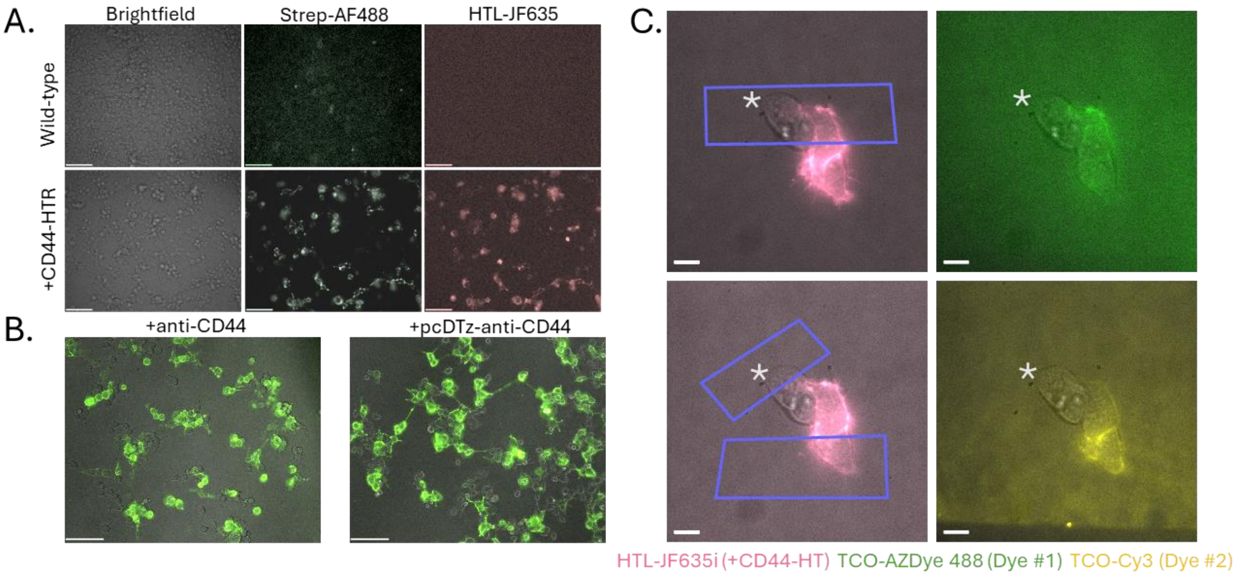


**Figure S3**. **Control experiments for antibody mediated photo-click.** A. Top, untransfected, bottom, CD44-HaloTag-expressing HEK cells, stained with biotinylated anti-CD44 and streptavidin-AF488 and HTL-JF635. Left: brightfield images, middle: AF488 fluorescence (green), right: JF635 fluorescence (red). Scale bar 100 µm. B. Merged brightfield and fluorescence images of CD44-HaloTag-expressing HEK cells. Left: stained with biotinylated antiCD44 and streptavidin-AF488 (green); right: stained with pcDTz-modified biotinylated antiCD44 and streptavidin-AF488 (green). Scale bar 100 µm. C. HEK cells transfected with CD44-HaloTag, treated with antiCD44-pcDTz and photopatterned. Purple indicates regions targeted with 405 nm light. Left image shows fluorescence from HaloTag ligand-JF635 (pink), indicating expression of the construct, overlayed on brightfield images. Right images show fluorescence of TCO-dye. Top: first uncaging and perfusion with TCO-AZdye488 (green). Bottom: second uncaging and perfusion with TCO-Cy3 (yellow). Scale bar 10 µm. Asterisk indicates untransfected cell.

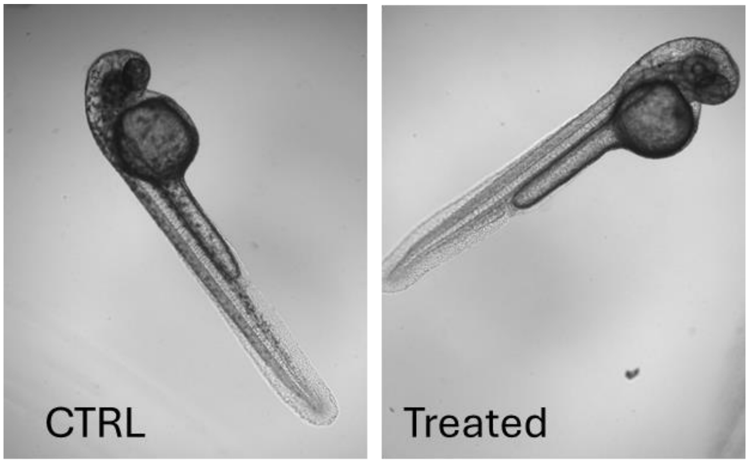


**Figure S4. Fish develop normally after photo-patterning.** Representative images of zebrafish, taken at 60 hours post fertilization, after control (fish water rinses, left) or photo-click surface patterning (right) at 14 hpf.

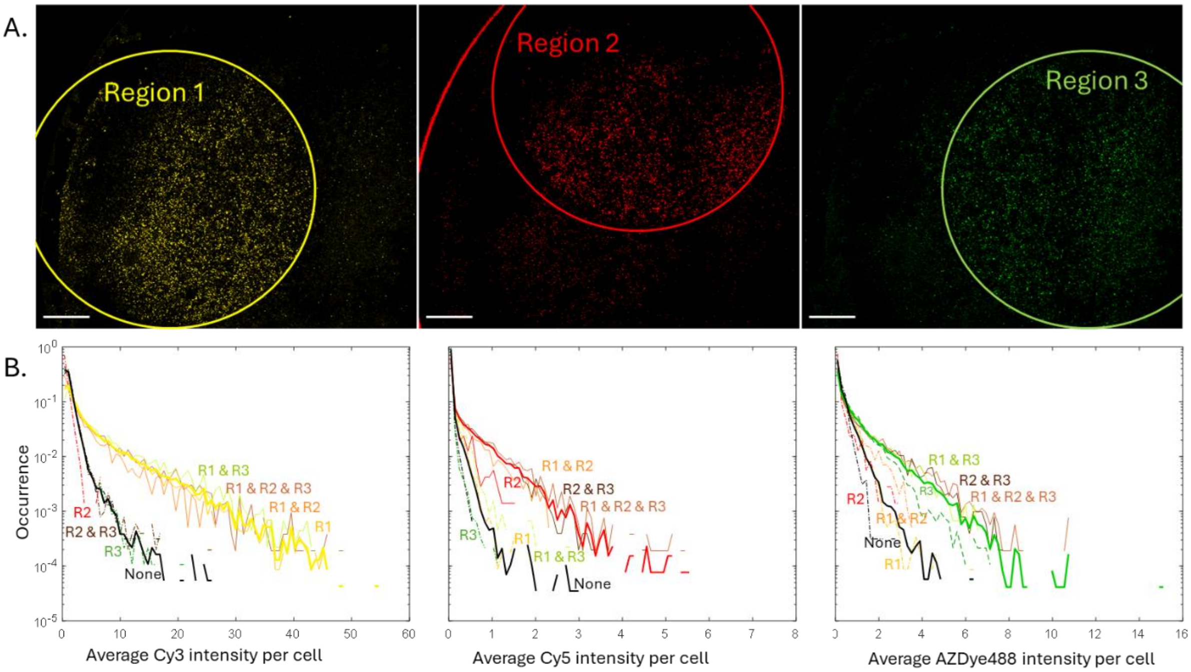


**Figure S5. Combinatorial photo-labeling of HTR-PDGFR-expressing HEK cells, analyzed by image processing.** A. Images of stained cells in each color channel separately (left: TCO-Cy3, yellow; middle: TCO-Cy5, red; right: TCO-AZDye488, green). Scale bar 1 mm. B. Frequency of cell intensities. Thick colored lines (left: yellow, middle: red, right: green) show cell intensities inside all regions illuminated at indicated stage, thick black line shows intensities for cells not illuminated at that stage. Thin lines denote all possible overlap regions, with solid lines indicating regions expected to retain dye, and dashed lines indicating those expected to be dark.

**Details of preparation and characterization of novel pcDTz compounds**

Common materials or chemical reagents were purchased commercially and used without further purification. All reactions were monitored by thin-layer chromatography (TLC), or high-resolution mass spectra (HR-MS). TLC was performed using silica gel plate (GF254, 0.23 mm) which were visualized with a UV lamp (254 nm and 365 nm). HR-MS were measured with an Agilent 6530 Accurate-Mass Quadrupole Time-of-Flight (Q-TOF) LC/MS system equipped with electrospray ionization (ESI). Nuclear magnetic resonance (NMR) spectra were recorded on a Quantum-I 400 or AVANCE NEO 700 spectrometer with TMS as the internal standard. Chemical shifts (δ) were reported in parts per million (ppm) relative to residual solvent peaks. Column chromatography was carried out using Biotage Rening Cartridge (particle size 0.040−0.063 mm) using technical grade solvents.

**Scheme 1**

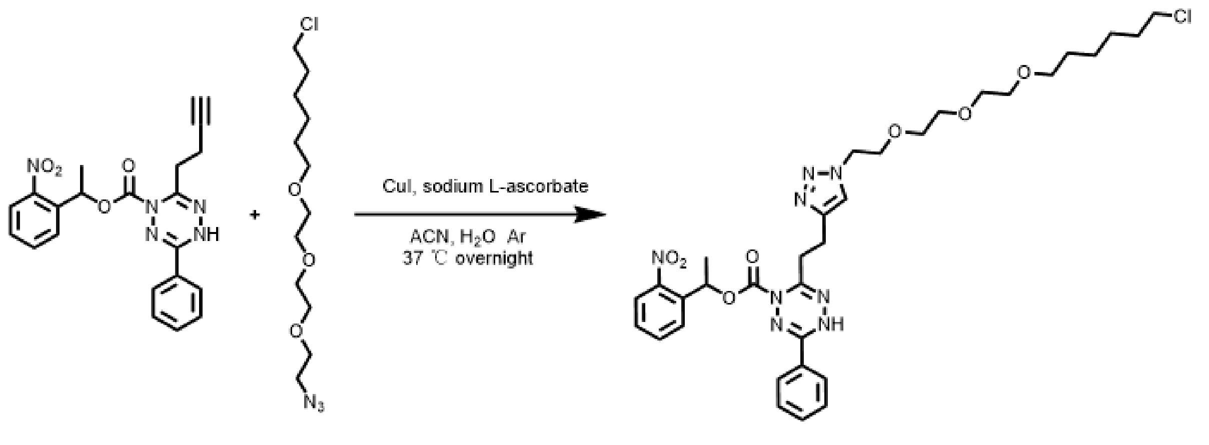


S1 halotag-azide HTL-pcDTz

Preparation of 1-(2-nitrophenyl)ethyl6-(2-(1-(2-(2-(2-((6chlorohexyl)oxy)ethoxy)ethoxy)ethyl)-1H-1,2,3-triazol-4-yl)ethyl)-3-phenyl-1,2,4,5-tetrazine-1(4*H*)-carboxylate (**HTL-pcDTz**):

1-(2-nitrophenyl)ethyl 6-(but-3-yn-1-yl)-3-phenyl-1,2,4,5-tetrazine-1(4*H*)-carboxylate (compound **S1**) was prepared according to the reported literature [1]. Under argon, 12 mL of $CH_3CN/H_2O$ (v/v = 2/1) was added to a mixture of photocaged dihydrotetrazine compound **S1** (127 mg, 0.31 mmol), 1-(2-(2-(2-azidoethoxy)ethoxy)ethoxy)-6-chlorohexane (125 mg, 0.42 mmol), CuI (106 mg, 0.56 mmol), and sodium L-ascorbate (110 mg, 0.56 mmol) at room temperature. The reaction mixture was stirred at 37 °C overnight. Upon completion, the reaction solvent was removed under reduced pressure. The residue was purified by column chromatography (4% MeOH/ $CH_2Cl_2$), and the desired product **HLT-pcDTz** was obtained (71 mg, 32% yield). **$^1$H NMR** (700 MHz, Chloroform-*d*) δ 8.02 (s, 1H), 7.98 (d, *J* = 8.2 Hz, 1H), 7.82 – 7.78 (m, 3H), 7.65 (t, *J* = 7.6 Hz, 1H), 7.53 (q, *J* = 8.3, 7.8 Hz, 1H), 7.48 (d, *J* = 7.6 Hz, 2H), 7.43 (t, *J* = 7.8 Hz, 1H),

6.46 (q, $J$ = 6.5 Hz, 1H), 4.44 (t, $J$ = 5.0 Hz, 2H), 3.77 (t, $J$ = 5.0 Hz, 2H), 3.61 (dd, $J$ = 5.9, 3.1 Hz, 2H), 3.59 – 3.54 (m, 4H), 3.50 (dd, $J$ = 8.1, 5.5 Hz, 4H), 3.42 (t, $J$ = 6.7 Hz, 2H), 3.08 (s, 2H), 2.98 (s, 2H), 1.77 (d, $J$ = 6.6 Hz, 3H), 1.73 (p, $J$ = 6.8 Hz, 2H), 1.54 (p, $J$ = 6.8 Hz, 2H), 1.43 – 1.37 (m, 2H), 1.32 (tt, $J$ = 9.9, 5.7 Hz, 2H). **$^{13}$C NMR** (176 MHz, $CDCl_3$) δ 155.47, 149.98, 147.45, 146.21, 143.76, 138.12, 133.89, 131.74, 128.90, 128.84, 128.43, 127.40, 126.76, 124.56, 122.62, 71.26, 70.61, 70.54, 70.49, 70.42, 70.04, 69.44, 53.72, 50.16, 45.06, 32.49, 30.91, 29.36, 26.64, 25.37, 22.33, 22.25. **HR-MS**: *m/z* = 699.30103 $[M+H]^+$, calc'd for $C_{33}H_{44}O_7N_8Cl$: 699.30160.

**Scheme 2**

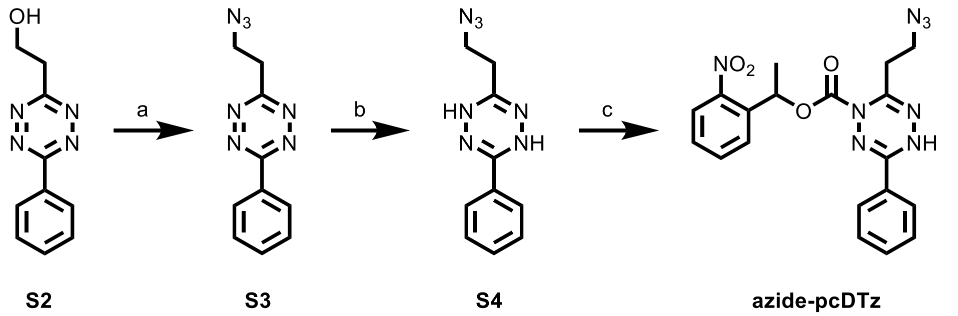


Reagents and conditions: a) compound **S2** (1 eq), MsCl (3 eq), triethylamine (3 eq), DMAP (1 eq), $CH_2Cl_2$, 0 °C to r.t., under argon, then $NaN_3$ (5 eq), DMF, yield 40%; b) compound **S3** (1 eq), Thiourea dioxide (2 eq), DMF/$H_2O$ (v/v = 1/2), 95 °C, yield: 74%; c) compound **S4** (1 eq), 1-(2-nitrophenyl)ethyl carbonochloridate (1.5 eq), Tol /Py (v/v =1/5) , 37 °C, under argon, yield: 83%.

**Preparation of 3-(2-azidoethyl)-6-phenyl-1,2,4,5-tetrazine (compound S3)**

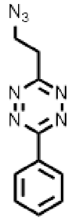

2-(6-phenyl-1,2,4,5-tetrazin-3-yl)ethan-1-ol **(compound S2)** was prepared according to the reported literature [2]. Under argon, triethylamine (617 µL, 4.5 mmol), MsCl (343 µL 4.5 mmol) and DMAP (182 mg, 1.5 mmol) were added to a solution of 2-(6-phenyl-1,2,4,5-tetrazin-3-yl)ethan-1-ol (300 mg, 1.5 mmol) in $CH_2Cl_2$ (8 mL) at room temperature. Upon completion, the reaction mixture was concentrated under reduced pressure. The residue was purified by column chromatography ($CH_2Cl_2$ as the eluent), and the 2-(6-phenyl-1,2,4,5-tetrazin-3-yl)ethyl methanesulfonate was obtained. Then, $NaN_3$ (309 mg, 4.75 mmol) was added to a solution of 2-(6-phenyl-1,2,4,5-tetrazin-3-yl)ethyl methanesulfonate in DMF (6 mL), the mixture stirred at room temperature under argon for 6 h. Then, the reaction mixture was extracted with $CH_2Cl_2$ and washed with a saturated solution of NaCl in water. The extract was combined, dried over $Na_2SO_4$, and concentrated under reduced pressure. The residue was purified by column chromatography (50% Heaxane/ EtOAc), and the desired product S3 was obtained (132 mg, 40% yield). **$^1$H NMR** (400 MHz, Chloroform-*d*) δ 8.65 – 8.58 (m, 2H), 7.69 – 7.56 (m, 3H), 4.03 (t, $J$ = 6.7 Hz, 2H), 3.64 (t, $J$ = 6.7 Hz, 2H). **$^{13}$C NMR** (101 MHz, CHLOROFORM-*D*) δ 167.26, 164.73, 132.97, 131.63, 129.41, 128.22, 48.84, 34.54. **HR-MS**: *m/z* = 228.09908 $[M+H]^+$, calc'd for $C_{10}H_{10}N_7$: 228.09922.

**Preparation of 3-(2-azidoethyl)-6-phenyl-1,4-dihydro-1,2,4,5-tetrazine (compound S4)**

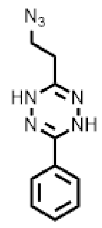


Under argon, the thiourea dioxide (76 mg, 0.7 mmol) was added to a solution of **S3** (80 mg, 0.35 mmol) in 15 mL of DMF/$H_2O$ (v/v = 1/2) at room temperature. The reaction mixture was stirred at 95 °C for 1–2 hours. Upon completion, the color of the reaction mixture changed from pink to light yellow. Under argon, 50 mL of EtOAc was added the reaction mixture, which was washed by 30 mL $H_2O$. The extract was combined and concentrated by reduced pressure. The residue was purified by column chromatography ($CH_2Cl_2$ as the eluent), and the desired product compound **S4** was obtained (60 mg, 74% yield). **$^1$H NMR** (400 MHz, Chloroform-*d*) δ 7.62 (dq, $J$ = 6.6, 1.4 Hz, 1H), 7.43 (dddd, $J$ = 11.7, 8.9, 5.4, 1.5 Hz, 1H), 3.66 – 3.58 (m, 1H), 2.47 (td, $J$ = 6.6, 1.1 Hz, 1H). **$^{13}$C NMR** (101 MHz, CHLOROFORM-*D*) δ 148.56, 148.35, 130.85, 130.19, 128.99, 126.07, 48.04, 30.36. **HR-MS**: *m/z* = 230.11452 $[M+H]^+$, calc'd for $C_{10}H_{12}N_7$: 230.11487.

**Preparation of 1-(2-nitrophenyl)ethyl 6-(2-azidoethyl)-3-phenyl-1,2,4,5-tetrazine-1(4*H*)-carboxylate (azide-pcDTz)**

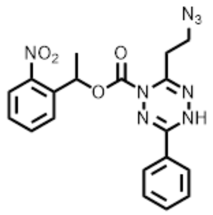


Under argon, the solution of **1-(2-nitrophenyl)ethyl carbonochloridate** (158 mg, 0.65 mmol) in toluene (1 mL) was added to a solution of compound **S4** (60 mg, 0.26 mmol) in pyridine (5 ml) at 0 ℃. The reaction mixture was stirred at 37 °C for 24 h. Upon completion, the reaction solvent was removed under reduced pressure. The residue was purified by column chromatography (1% MeOH/ $CH_2Cl_2$), and the azide-pcDTz was obtained (70 mg, 83% yield). **$^1$H NMR** (700 MHz, Chloroform-*d*) δ 8.00 (dd, $J$ = 8.2, 1.4 Hz, 1H), 7.80 (dd, $J$ = 7.9, 1.5 Hz, 1H), 7.76 – 7.74 (m, 2H), 7.66 (td, $J$ = 7.6, 1.5 Hz, 1H), 7.58 – 7.55 (m, 1H), 7.50 (dd, $J$ = 8.4, 6.9 Hz, 2H), 7.48 – 7.44 (m, 2H), 3.59 – 3.39 (m, 2H), 2.98 (td, $J$ = 6.5, 1.9 Hz, 2H), 1.78 (d, $J$ = 6.6 Hz, 3H). **$^{13}$C NMR** (176 MHz, $CDCl_3$) δ 154.95, 150.05, 147.47, 142.25, 138.02, 133.87, 131.96, 129.10, 128.66, 128.52, 127.31, 126.57, 124.67, 70.76, 47.78, 31.01, 22.26. **HR-MS**: *m/z* = 423.15198 $[M+H]^+$, calc'd for $C_{19}H_{19}O_4N_8$: 423.15238.

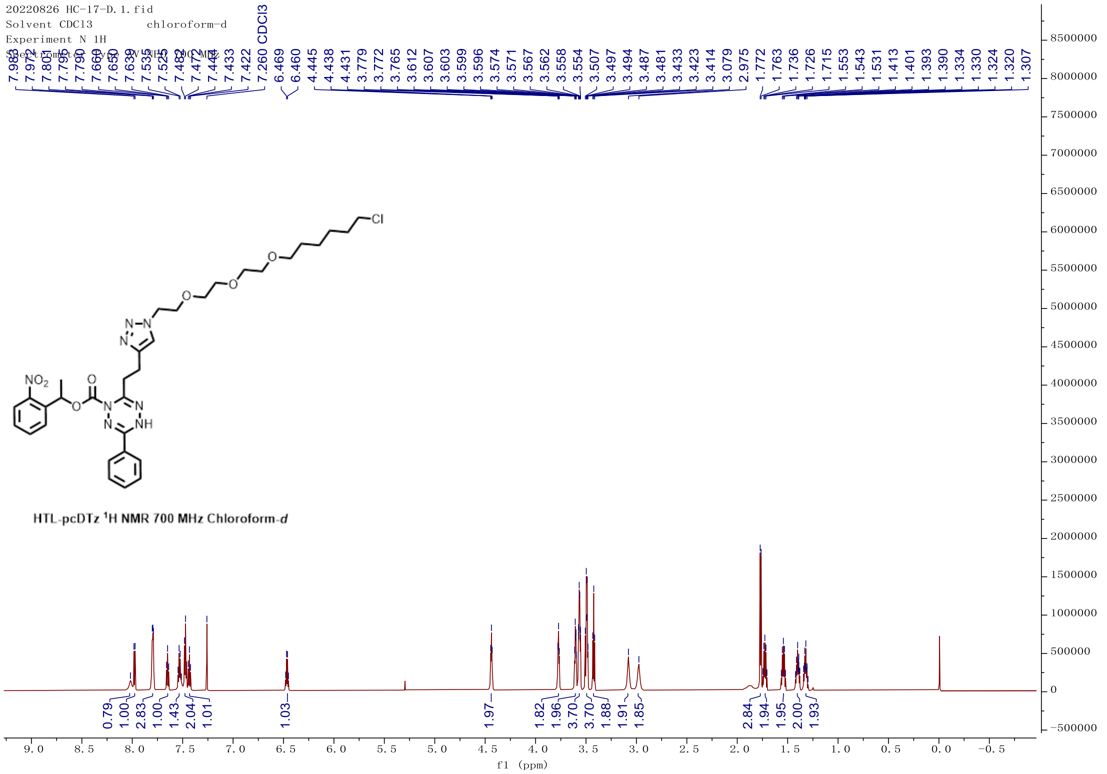
20220826 HC-17-D.1.fid
Solvent CDCl3 chloroform-d
Experiment N 1H
7.260 CDCl3
7.972 7.801 7.795 7.790 7.660 7.650 7.639 7.535 7.525 7.482 7.472 7.444 7.433 7.422
6.469 6.460 4.445 4.438 4.431 3.779 3.772 3.765 3.612 3.607 3.603 3.599 3.596 3.574 3.571 3.567 3.562 3.558 3.554 3.507 3.497 3.494 3.487 3.481 3.433 3.423 3.414 3.079 2.975 1.772 1.763 1.736 1.726 1.715 1.553 1.543 1.531 1.413 1.401 1.393 1.390 1.334 1.330 1.324 1.320 1.307
NO2
O
N
NH
Cl
HTL-pcDTz 1H NMR 700 MHz Chloroform-d
0.79 1.00 2.83 1.00 1.43 2.04 1.01 1.03 1.97 1.82 1.96 3.70 3.70 1.88 1.91 1.85 2.84 1.94 1.95 2.00 1.93
f1 (ppm)
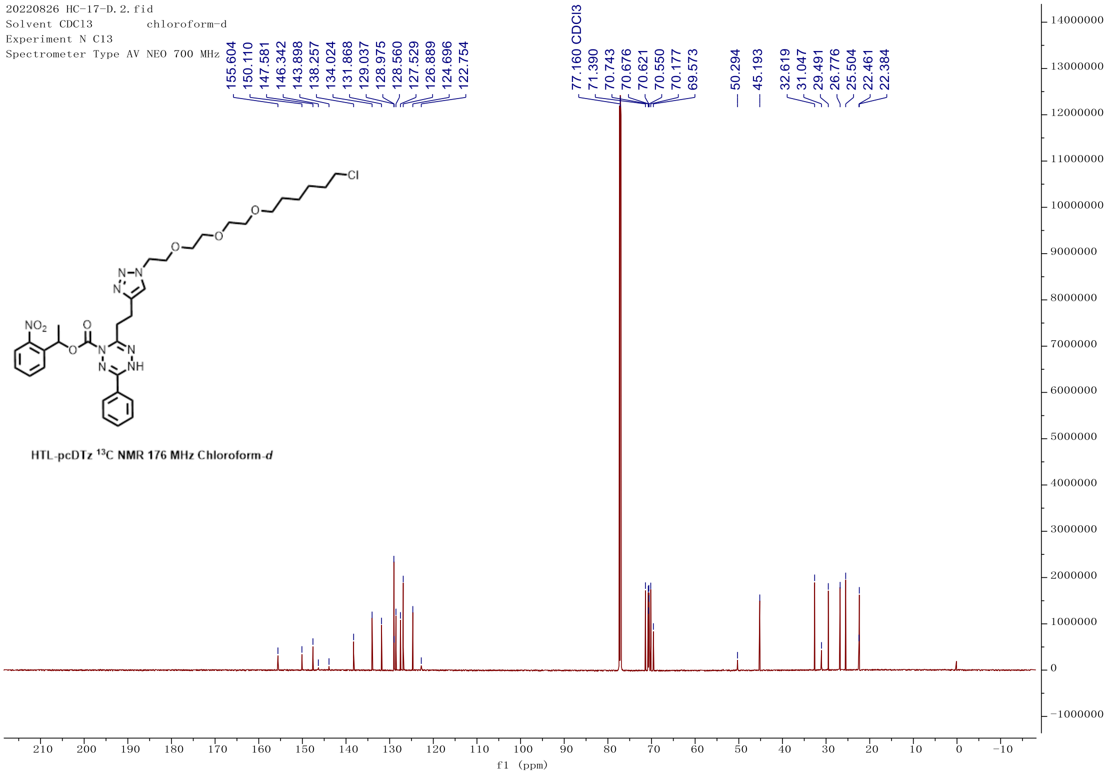
20220826 HC-17-D.2.fid
Solvent CDCl3 chloroform-d
Experiment N C13
Spectrometer Type AV NEO 700 MHz
155.604 150.110 147.581 146.342 143.898 138.257 134.024 131.868 129.037 128.975 128.560 127.529 126.889 124.696 122.754
77.160 CDCl3 71.390 70.743 70.676 70.621 70.550 70.177 69.573
50.294 45.193
32.619 31.047 29.491 26.776 25.504 22.461 22.384
NO2
O
N
NH
Cl
HTL-pcDTz 13C NMR 176 MHz Chloroform-d
f1 (ppm)

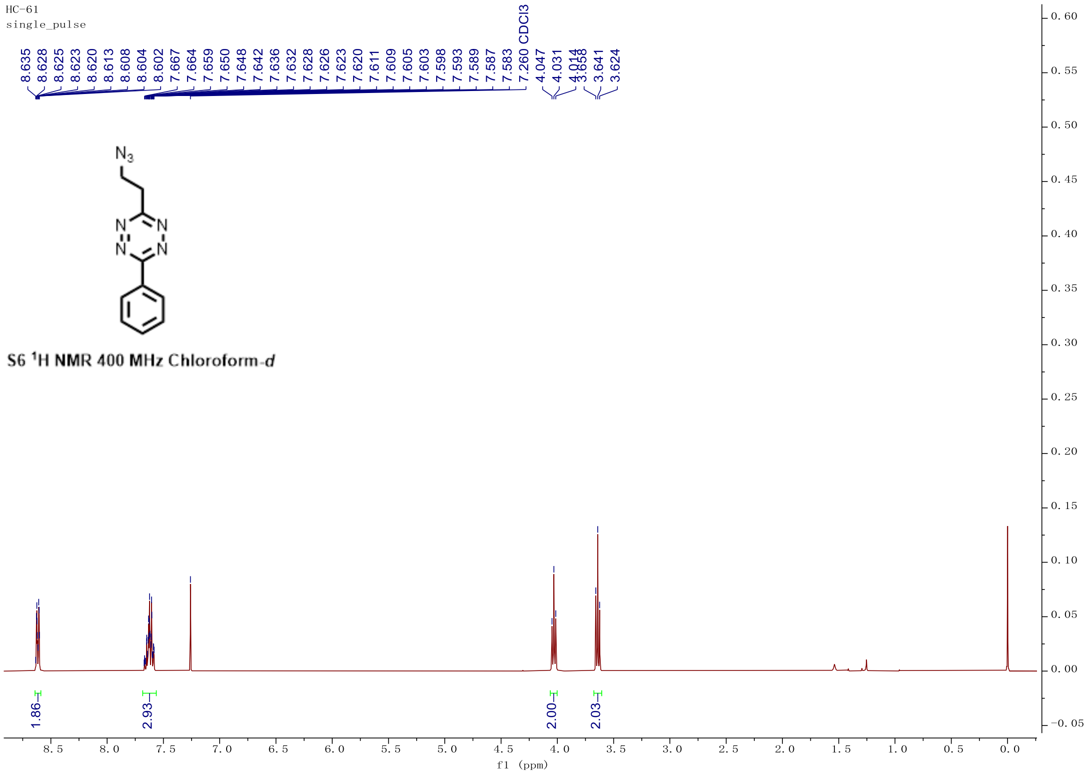
HC-61
single_pulse
8.635 8.628 8.625 8.623 8.620 8.613 8.608 8.604 8.602 7.667 7.664 7.659 7.650 7.648 7.642 7.636 7.632 7.628 7.626 7.623 7.620 7.611 7.609 7.605 7.603 7.598 7.593 7.589 7.587 7.583 7.260 CDCl3 4.047 4.031 4.014 3.658 3.641 3.624
N3
N N N N
S6 1H NMR 400 MHz Chloroform-d
1.86
2.93
2.00
2.03
f1 (ppm)

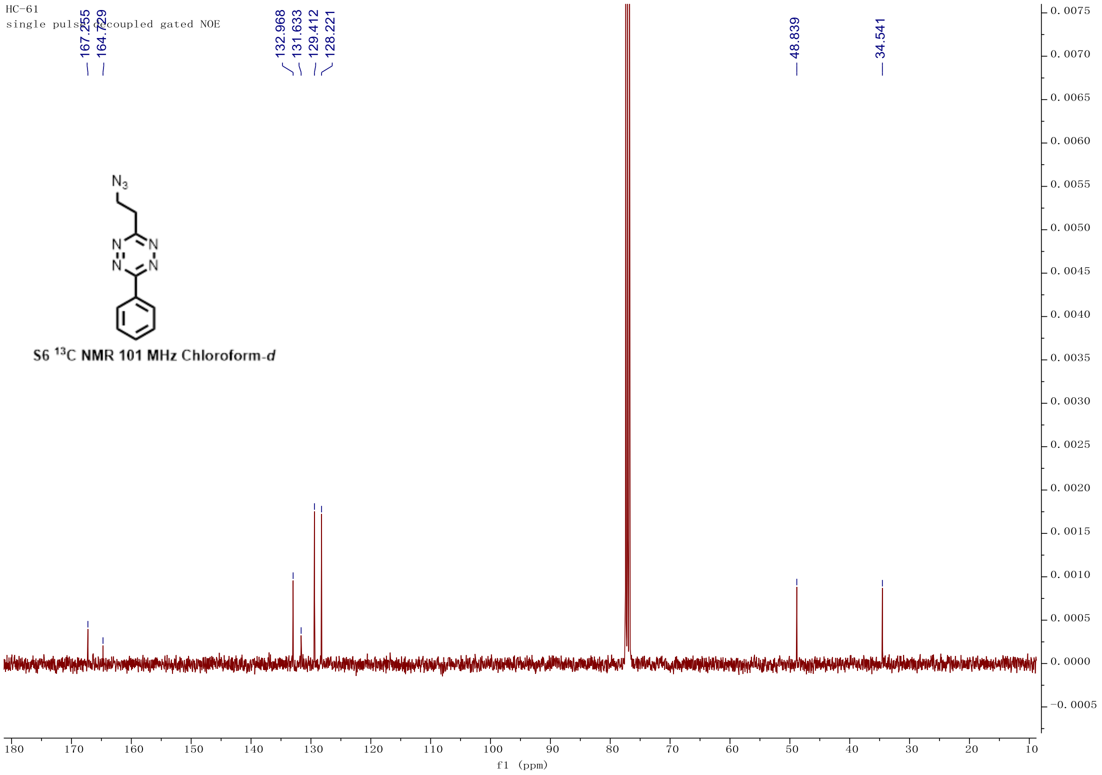
HC-61
167.255 164.729 132.968 131.633 129.412 128.221 48.839 34.541
N3
N N N N
S6 13C NMR 101 MHz Chloroform-d
f1 (ppm)

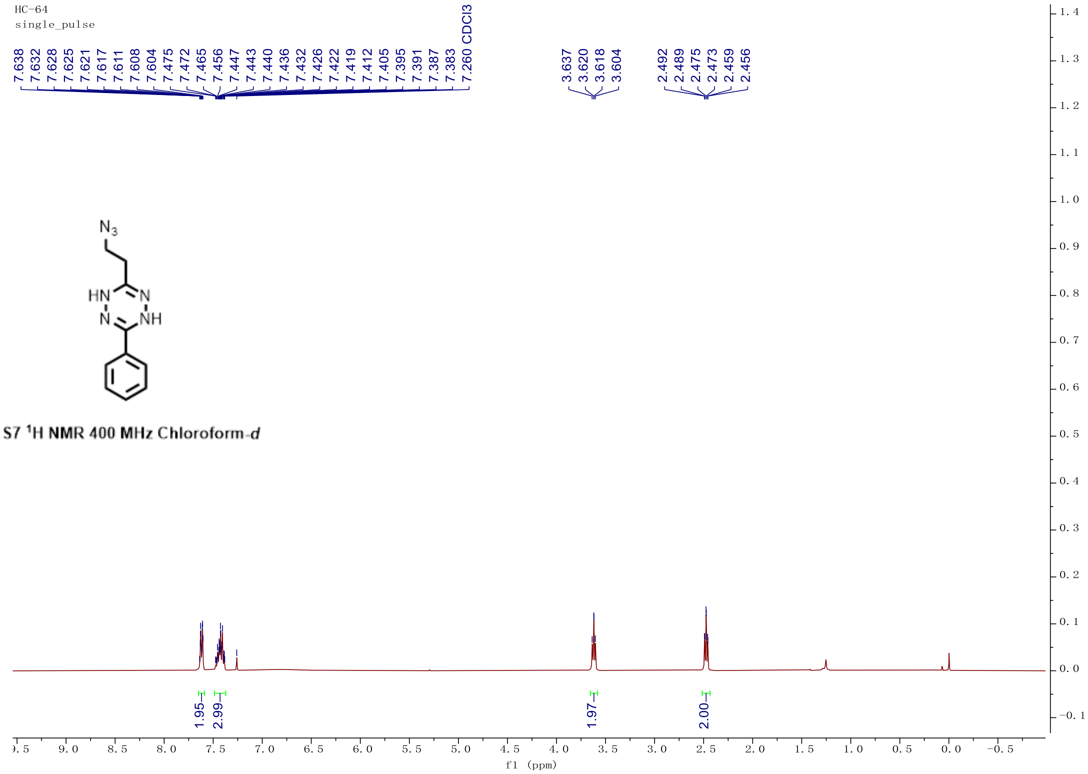
HC-64
single_pulse
7.638
7.632
7.628
7.625
7.621
7.617
7.611
7.608
7.604
7.475
7.472
7.465
7.456
7.447
7.443
7.440
7.436
7.432
7.426
7.422
7.419
7.412
7.405
7.395
7.391
7.387
7.383
7.260 CDCl3
3.637
3.620
3.618
3.604
2.492
2.489
2.475
2.473
2.459
2.456
N3
HN
N
N
NH
S7 1H NMR 400 MHz Chloroform-d
1.95
2.99
1.97
2.00
f1 (ppm)

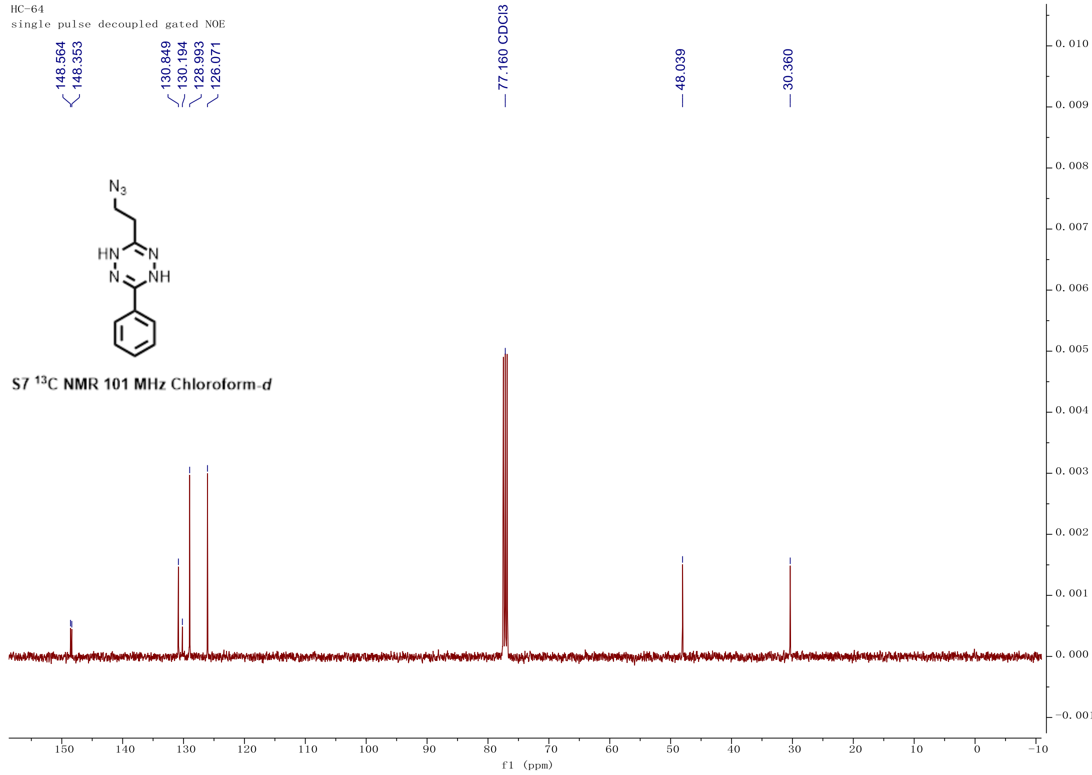
HC-64
single pulse decoupled gated NOE
148.564
148.353
130.849
130.194
128.993
126.071
77.160 CDCl3
48.039
30.360
N3
HN
N
N
NH
S7 13C NMR 101 MHz Chloroform-d
f1 (ppm)

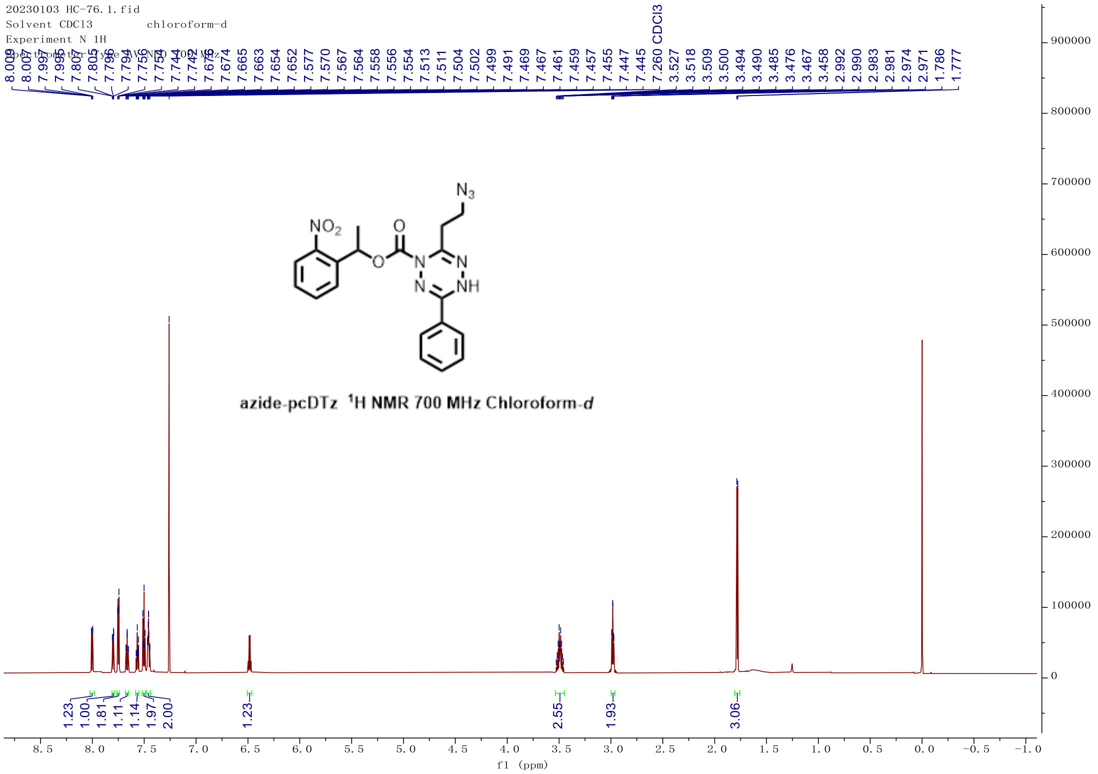
20230103 HC-76.1.fid
Solvent CDCl3 chloroform-d
Experiment N 1H
azide-pcDTz 1H NMR 700 MHz Chloroform-d
f1 (ppm)

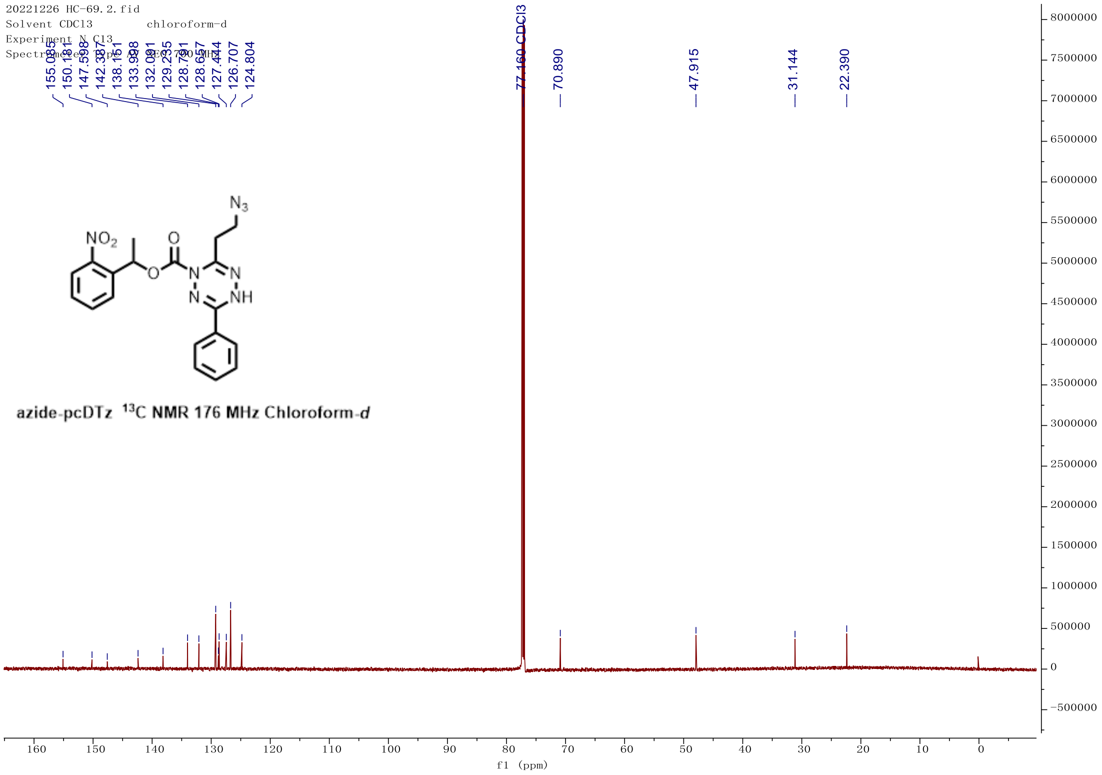
20221226 HC-69.2.fid
Solvent CDCl3 chloroform-d
Experiment N C13
azide-pcDTz 13C NMR 176 MHz Chloroform-d
f1 (ppm)

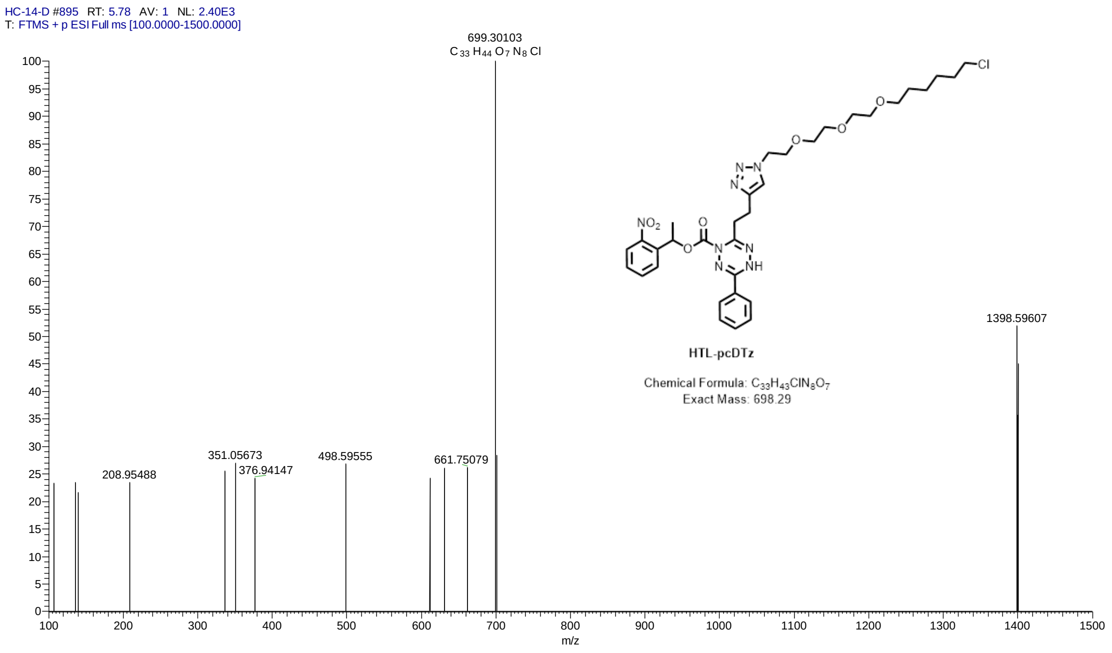

HC-14-D #895 RT: 5.78 AV: 1 NL: 2.40E3
T: FTMS + p ESI Full ms [100.0000-1500.0000]
699.30103
C 33 H 44 O 7 N 8 Cl
1398.59607
208.95488
351.05673
376.94147
498.59555
661.75079
m/z
HTL-pcDTz
Chemical Formula: C33H43ClN8O7
Exact Mass: 698.29


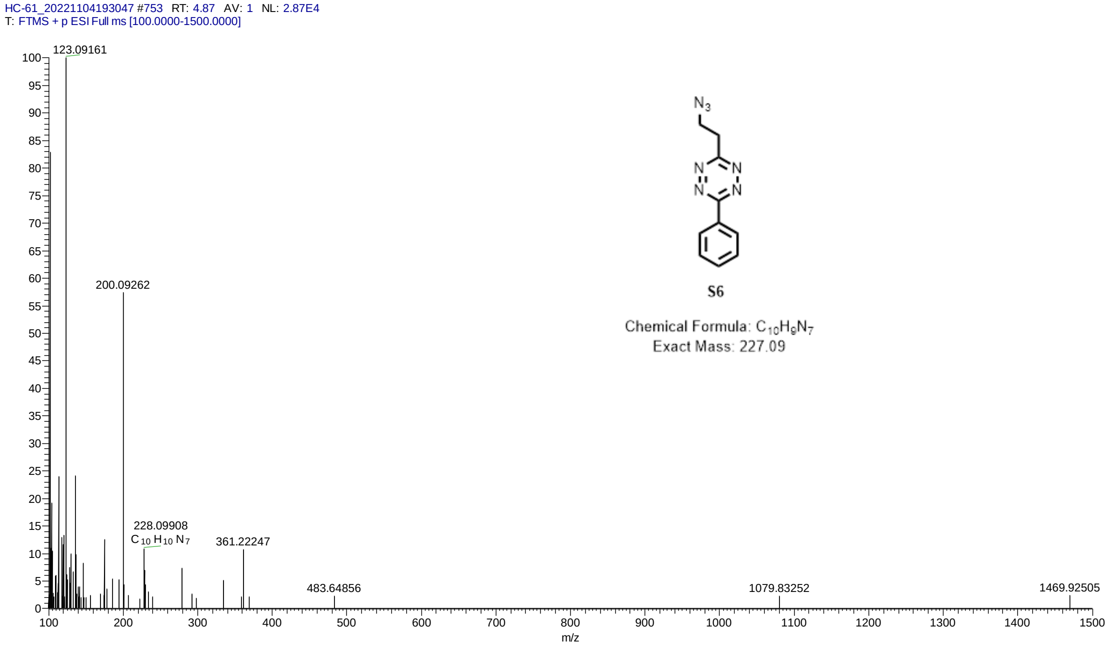

HC-61_20221104193047 #753 RT: 4.87 AV: 1 NL: 2.87E4
T: FTMS + p ESI Full ms [100.0000-1500.0000]
123.09161
200.09262
228.09908
C 10 H 10 N 7
361.22247
483.64856
1079.83252
1469.92505
m/z
S6
Chemical Formula: C10H9N7
Exact Mass: 227.09

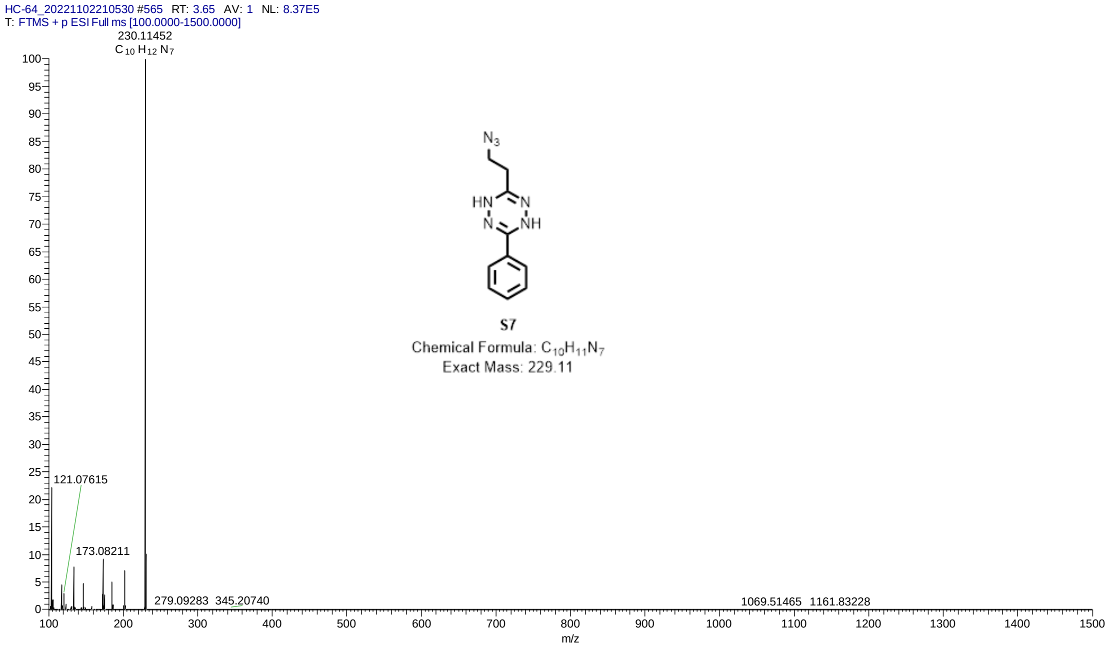


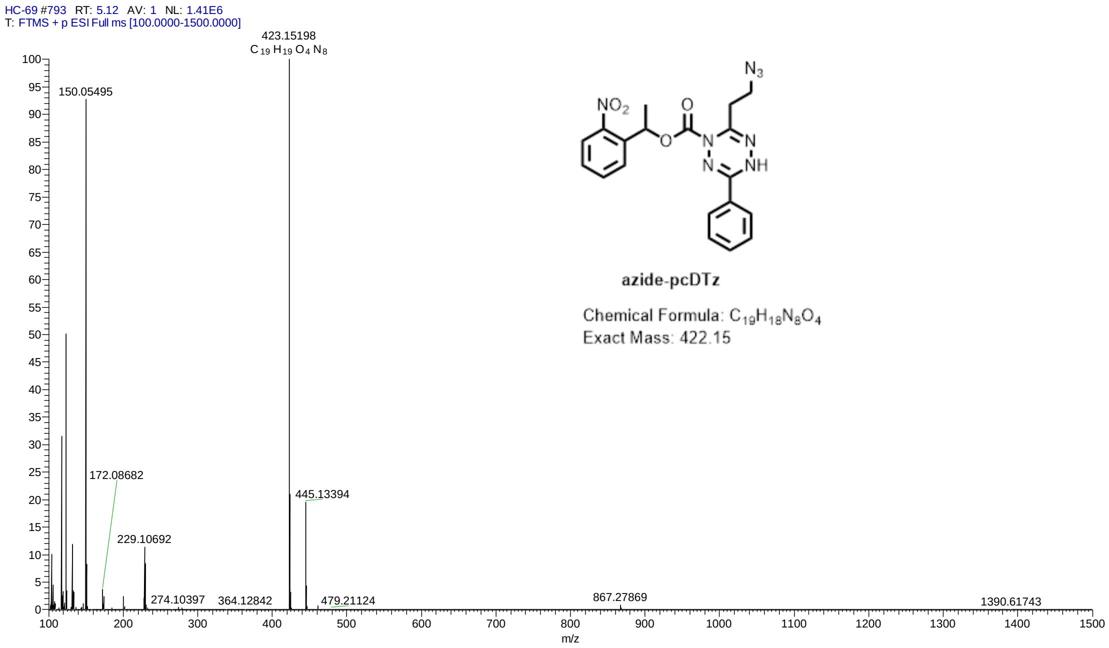


### References:


[1] Liu L, Zhang D, Johnson M, et al. Light-activated tetrazines enable precision live-cell bioorthogonal chemistry [J]. Nat Chem, 2022, 14: 1078.

[2] Mao W, Shi W, Li J, et al. Organocatalytic and Scalable Syntheses of Unsymmetrical 1,2,4,5-Tetrazines by Thiol-Containing Promotors [J]. Angewandte Chemie International Edition, 2018, 58(4): 1106-9.
